# Analysis and design of single-cell experiments to harvest fluctuation information while rejecting measurement noise

**DOI:** 10.1101/2021.05.11.443611

**Authors:** Huy D. Vo, Linda Forero, Luis Aguilera, Brian Munsky

## Abstract

Despite continued technological improvements, measurement errors will always reduce or distort the information that any real experiment can provide to quantify cellular dynamics. This problem becomes even more serious in the context of cell signaling studies that are specifically designed to quantify heterogeneity in single-cell gene regulation, where important RNA and protein copy numbers are themselves subject to the inherently random fluctuations of biochemical reactions. It is not clear how measurement noise should be managed in addition to other experiment design variables (e.g., sampling size, measurement times, or perturbation levels) to ensure that collected data will provide useful insights on signaling or gene expression mechanisms of interest. To address these fundamental single-cell analysis and experiment design challenges, we propose a computational framework that takes explicit consideration of measurement errors to analyze single-cell observations and Fisher Information Matrix-based criteria to decide between experiments. Using simulations and single-cell experiments for a reporter gene controlled by an HIV promoter construct, we demonstrate how our approach can analyze and redesign experiments to optimally harvest fluctuation information while mitigating the effects of image distortion.

## 1 Introduction

Heterogeneity in signaling and gene expression at the single-cell level has wide-ranging biological and clinical consequences, from bacterial persistence ([46, 26, 27, 16]) and viral infections ([78, 91, 68, 8]) to tumor heterogeneity ([7, 52]). Beside genetic and environmental factors, a significant degree of heterogeneity is caused by biochemical noise ([53, 66, 17, 3]). Therefore, even genetically identical cells grown in the same experimental conditions may display variability in their response to environmental stimuli. This variability, often termed *intrinsic noise* when it originates within the pathway of interest or *extrinsic noise* when it originates outside the pathway of interest, obscures the underlying mechanisms when viewed through the lens of deterministic models and bulk measurements ([57]). Yet, this so-called noise can be highly informative when examined through the lens of single-cell measurements coupled with the mathematical modeling framework of the Chemical Master Equation (CME) ([28, 1, 81]). Through this joint experimental and modeling approach, mechanisms of signaling and single-cell gene expression can be explained, predicted ([58, 59, 57]), or even controlled ([56, 55, 5, 22]).

In this paper, we are concerned with how and when stochastic models based on the CME could be inferred with high confidence from single-cell experiments such as flow cytometry or optical microscopy (e.g., single-molecule *in situ* hybridization, smFISH), from which data sets are produced in the form of fluorescence intensity histograms ([36, 48, 93, 50]) or molecular count histograms ([20, 64, 47]). These measurements, like any other experimental technique, can be corrupted by errors arising from imprecise detection or data processing methods. In flow cytometry, variations in the binding efficiency, binding specificity, or intensity of individual fluorescent reporters or probes, or the existence of background fluorescence, are unavoidable perturbations that may obscure the true copy number of RNA or protein ([58, 70, 86]). While smFISH data sets are generally considered to provide the’gold standard’ for measuring transcriptional heterogeneity, the estimation of molecular copy numbers in single cells depends heavily on image segmentation and spot counting algorithms ([4, 62, 65, 39]) that involve several threshold parameters, which are set *ad hoc* and typically vary from one expert to another. In addition, the sensitivity of smFISH measurements can be heavily affected by the length of the target mRNAs, the number of probes ([76]), or the number of hybridization steps ([92]).

The inescapable fact of measurement noise motivates intriguing questions on the design of single-cell experiments. For instance, under what conditions can detailed statistical measurement noise models and cheap single-cell measurements combine to replace accurate, yet more expensive experimental equipment? How approximate or coarse (and therefore fast) can image processing be while still faithfully retaining knowledge about parameters or mechanisms of interest? These questions are becoming more pressing as new advances in fluorescence tagging and microscopy technology are leading to more sophisticated experimental protocols to produce ever-increasing data on complex signaling and gene expression networks. Addressing these challenges is non-trivial, as it requires careful consideration of the potentially nonlinear combination of measurement noise and the biological question of interest.

The key contribution of this study is to provide new model-driven experiment analysis and design approaches (Fig 1) that include explicit consideration of probabilistic measurement errors in single-cell observations. The process begins with one or more hypotheses written in the form of stochastic gene regulation models (Fig 1A) with uncertain guesses for parameters or mechanisms. Predictions from these biophysical models are then coupled with empirically determined or physically estimated statistical models, known as *Probabilistic Distortion Operators* (PDOs), that explicitly estimate the effects of different types of measurement errors that could be temporal, discrete, non-symmetric, non-Gaussian, or even the result of their own stochastic process dynamics (Fig 1B). During model inference, the biophysical and measurement distortion models are simultaneously optimized to best explain the data (Fig 1C, right) and to quantify uncertainty in their respective parameters (Fig 1C, left). During experiment design (Fig 1D), sensitivity analysis is used to estimate which combinations of experimental conditions, sampling procedures, or measurement strategies can be expected to provide the most information to constrain the current set of hypotheses, and the procedure is iterated in subsequent rounds of experimentation and model refinement.

**Figure 1:**
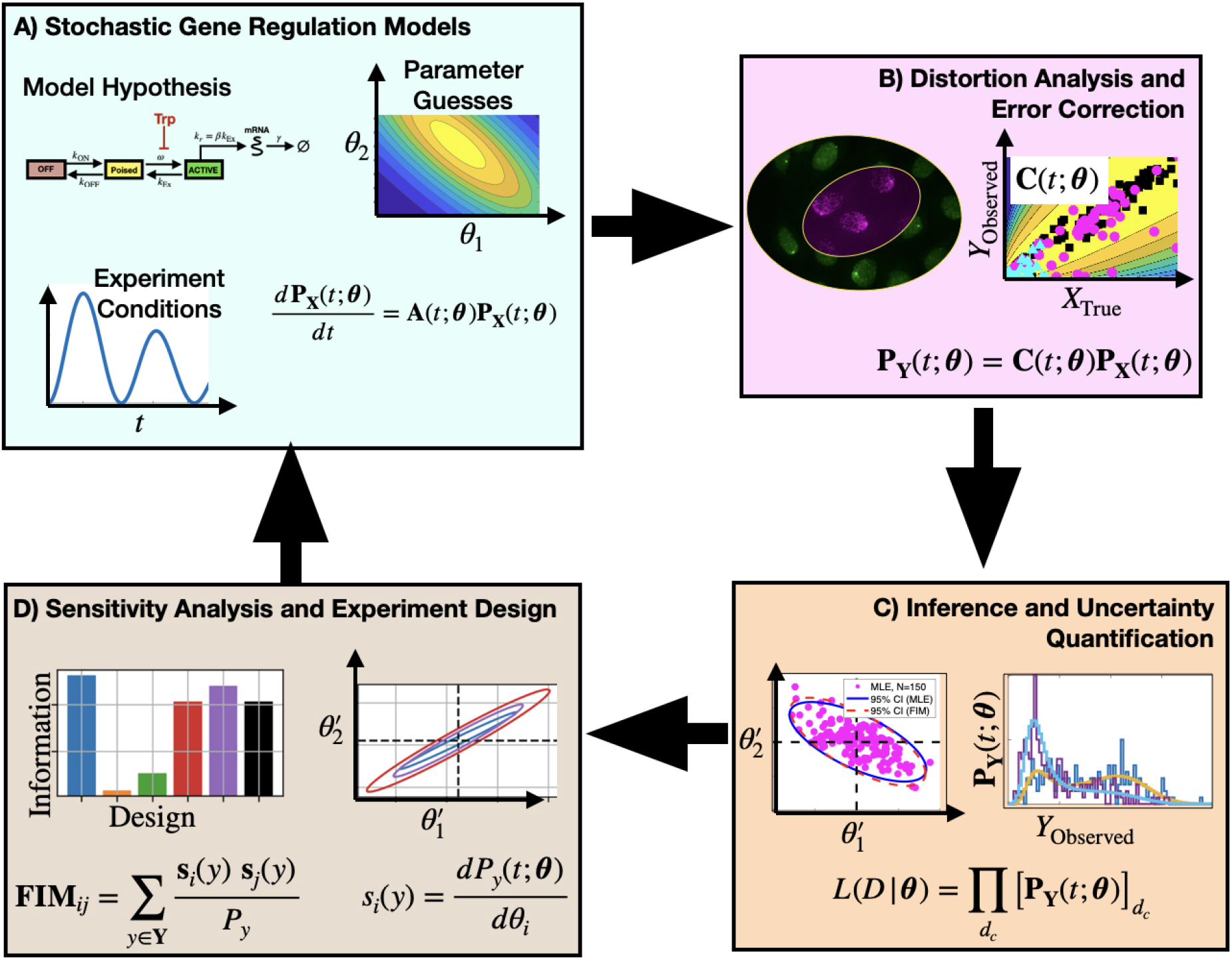
Proposed approach to analyze and design single-cell experiments under measurement distortion. **Box (A):** One or more mechanistic model hypotheses are proposed to describe gene regulation processes and are associated with prior parameter guesses. These models are combined with experiment design considerations (e.g., environmental, sampling, and measurement conditions) and the chemical master equation framework is used to predict statistics of single-cell gene expression. **Box B**: Empirical data and physical models are used to define a probabilistic distortion operator **C** to estimate how measurement errors (e.g., labeling inefficiencies, resolution limitations, or image processing errors) affect observations. **Box C**: Searches over model and PDO parameter space are conducted to identify model and parameter combinations that maximize the likelihood of the observed (i.e., distorted) data. **Box D**: Sensitivity analysis and the Fisher Information Matrix are computed and used to analyze the information content for different mechanistic models, distortion effects, or experiment designs. By estimating parameter uncertainties in the context of different forms of measurement distortion, subsequent experiment designs can help to alleviate measurement noise distortion effects (return to **Boxes A, B**, and **C**).

To address the specific question of model-driven single-cell experiment evaluation and design (Fig 1D), we adopt the framework of the Fisher Information Matrix (FIM). The FIM is the basis for a large set of tools for optimal experiment design in a myriad of science and engineering fields ([63, 18, 13, 69, 12, 89]), and it has been employed in the study of identifiability and robustness of deterministic ODEs models in system biology ([31, 87]) as well as for designing optimal bulk measurement experiments ([19, 25, 43]). As an early application of the FIM to stochastic modeling of gene expression, Komorowski et al. ([42]) devised a numerical method to compute the FIM based on the Linear Noise Approximation (LNA) of the CME, which they used to demonstrate the different impacts of time-series, time snapshots, and bulk measurements to parameter uncertainties. There have been subsequent work on approximating the FIM with moment closure techniques ([69, 70]), with demonstrable effectiveness on designing optimal optogenetic experiments([71]). Under an assumption of ideal measurements, Fox and Munsky ([23]) recently extended the Finite State Projection (FSP) algorithm ([23]) to compute a version of the FIM that allows for time-varying and non-linear models that result in discrete, asymmetric, and multi-modal single-cell expression distributions. By extending the FSP-based FIM ([23]) to also account for realistic measurement errors, our new approach can help scientists to decide which combinations of experimental conditions and measurement assays are best suited to reduce parameter uncertainties and differentiate between competing hypotheses.

To verify our proposed approaches, we simulate data for a simple bursting gene expression model under many different types of measurement errors, and we show that the FIM correctly estimates the effects that measurement distortions have on parameter estimation (we explore more complicated models and distortions in the Supplemental Text). To demonstrate the practical use of our approaches, we apply them to analyze single-cell data for the bursting and deactivation of a reporter gene controlled by an HIV promoter construct upon application of triptolide (Trp). We show that the iterative use of FSP to fit distorted experimental data, followed by FIM analysis to design subsequent experiments can lead to the efficient identification of a well-constrained model to explain and predict gene expression.

## 2. Methods

### 2.1 Stochastic modeling of gene expression

The expression dynamics of genes or groups of genes in single cells are often modeled by stochastic reaction networks ([54, 28, 1]). For these, the time-varying molecular copy numbers in single cells are treated as a Markov jump process*{****X***(*t*)*}*_*t≥*0_ whose sample paths ***x***(*t*) = (*x*_1_(*t*), …, *x*_*N*_ (*t*)) take place in discrete multi-dimensional space, where *x*_*i*_(*t*) is the integer count of species *i* at time *t*. Each jump in this process corresponds to the occurrence of one of the reaction events *R*_*k*_ (*k* = 1, …, *M*), which brings the cell from the state ***x***(*t*_−_) right before event time *t* to a new state ***x***(*t*_+_) = ***x***(*t*_−_) + ***v***_*k*_, where ***v***_*k*_ is the stoichiometry vector corresponding to the *k*-th reaction. The probabilistic rate at which each reaction occurs is characterized by its propensity (or reaction intensity) function, *α*_*k*_(*t*, ***x, θ***). The vector ***θ*** = (*θ*_1_, …, *θ*_*d*_) is a *d*-dimensional vector of *model parameters*. Intuitively, we interpret *α*_*k*_(*t*, ***x, θ***)Δ*t* as the probability for reaction *k* to fire during the waiting interval [*t, t* + Δ*t*) for a sufficiently small waiting time Δ*t*. The probability distributions ***p***_***X***_ (*t*, ***θ***) of single-cell gene product copy numbers model what is often termed *intrinsic noise* in gene expression. These distributions are the solution of the chemical master equation (CME)

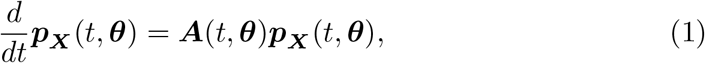

where ***A***(*t*, ***θ***) is the infinitesimal generator of the Markov process described above (see Supplemental Text S1 for detailed definition of **A**(*t*, ***θ***)). Extrinsic noise can be modeled by assuming a probabilistic variation for the model parameters, then integrate (1) over that distribution ([71]). However, we focus on intrinsic noise for the current investigation.

#### 2.1.1 Computing the likelihood of single-cell data

Consider a data set **D** that consists of *N*_*c*_ independent single-cell measurements (*t*_1_, ***c***_1_), …, 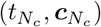 where ***c***_*i*_ is the vector of molecule counts of cell *i* measured at time *t*_*i*_. The likelihood function of the biophysical parameters ***θ*** given the data set **D** is given by

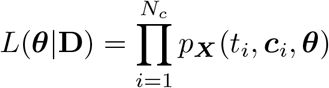

where *p*_***X***_ (*t*, ***x, θ***) is the probability of observing molecular counts ***x*** at time *t*, obtained by solving the CME (1). Taking the logarithm of both sides, we have the log-likelihood function

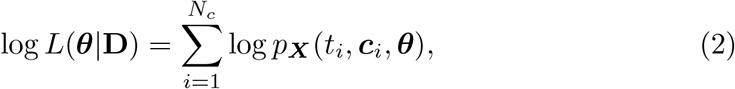

which is mathematically and numerically more convenient to work with.

#### 2.1.2 Sensitivity analysis

Taking the partial derivative of both sides of the CME (1) with respect to parameter *θ*_*l*_ (*l* = 1, …, *d*), we get

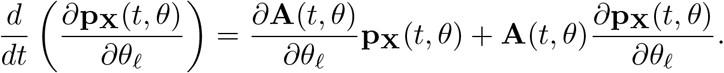

When the state space of the Markov process is finite, we can collect the equations above for *l* = 1, …, *d* along with the CME (1) to form a joint system involving the CME solution ***p***_***X***_ (*t, θ*) and its partial derivatives *s*_***X***,*l*_(*t*, ***θ***) ≡ *∂p*_***X***_ (*t*, ***θ***)*/∂θ*_*l*_, given by

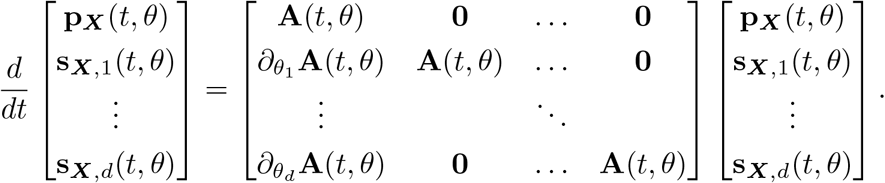

This forward sensitivity system can be solved numerically with any standard ODE solver. When the state space is infinite, a truncation algorithm based on extending the Finite State Projection ([23]) can be applied to approximately solve the forward sensitivity system (see SI Section 1 for more details).

Knowing the sensitivity of the distribution to parameter changes then allows us to compute the sensitivity of the log-likelihood function with respect to biophysical parameters *θ*_1_, …, *θ*_*d*_ using the formula

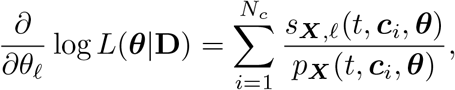

where *s*_***X***,*l*_(*t*, ***x, θ***) ≡ *∂p*_***X***_ (*t*, ***x, θ***)*/∂θ*_*l*_ is the sensitivity for the specific molecular counts ***x*** at time *t*.

#### 2.1.3 Modeling distortion of measurements

Let ***y***(*t*) be the multivariate measurement made on a single cell at time *t*, such as the discrete number of spots in an smFISH experiment, or the total fluorescence intensity, such as from a flow cytometry experiment. Because of random measurement noise, ***y***(*t*) is the realization of a random vector ***Y*** (*t*) that is the result of a random distortion of the true process ***X***(*t*). The probability mass (density) vector (function) ***p***_***Y***_ (*t*) of the discrete (continuous) observable measurement ***Y*** (*t*) is related to that of the true copy number distribution via a linear transformation of the form

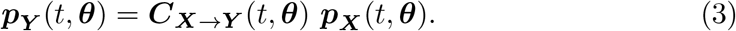

Mathematically, ***C***_***X***→***Y***_ (*t*, ***θ***) functions as a Markov kernel, and we shall call it the *Probabilistic Distortion Operator* (PDO) to emphasize the context in which it arises. Considered as a matrix whose rows are indexed by all possible observations ***y*** and whose columns are indexed by the CME states ***x***, it is given entry-wise as

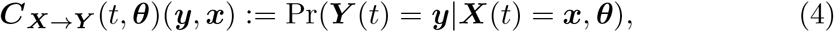

where Pr stands for probability mass if ***Y*** is discrete or probability density if ***Y*** is continuous. Together, equations (1) and (3) describe single-cell measurements as the time-varying outputs of a linear dynamical system on the space of probability distributions on the lattice of *N* -dimensional discrete copy number vectors. The output matrix of this dynamical system is the PDO ***C***_***X***→***Y***_. If the observations ***c***_1_, …, 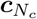 in the dataset ***D*** are assumed to be distorted according to the PDO ***C***, then the log-likelihood function (2) is changed into

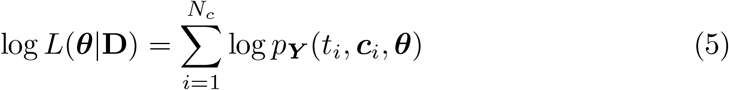

where *p*_***Y***_ (*t*, ***y, θ***) are point-wise probabilities of the distribution ***p***_***Y***_ (*t*, ***θ***) defined in (3).

There are many different ways to specify the PDO, depending on the specifics of the measurement method one wishes to model. In this paper, we demonstrate various examples where the PDO is formulated as probabilistic models that use simple distributions as building blocks (see Results and SI Section S3.2), or as deterministic binning/aggregation (Results and SI Section S3.2). One could even use a secondary CME to model the uncertain chemical kinetics of the measurement process, such as the random time needed to achieve chemical fixation in smFISH experiments or the dropout and amplification of mRNA that occurs during single-cell RNAseq experiments (SI Section S3.3), or distribution convolution to describe cell segmentation noise (SI Section S3.4). Despite the variety of ways these measurement noise models can be derived, they all lead to the same mathematical object (i.e., the PDO), which allows computation of the FIM associated with that particular noisy single-cell observation approach, and the effects of all PDO can be analyzed using the same computational procedure as we describe next.

#### 2.1.4 Computation of the Fisher Information Matrix for distorted experimental measurements

In practice, when closed-form solutions to the CME do not exist, a forward sensitivity analysis using an extension of the finite state projection algorithm can be used to evaluate the probability distribution ***p***_***X***_ (*t*, ***θ***) and its partial derivatives 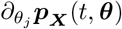 with respect to the kinetic parameters ([23]. Using eq. (3), we can transform these into the distribution ***p***_***Y***_ (*t*, ***θ***) of ***Y*** (*t*). Furthermore, the sensitivity indices 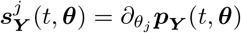, of the observable ***Y*** are computable by back-propagating the sensitivities of the noise-free measurement distributions through the PDO,

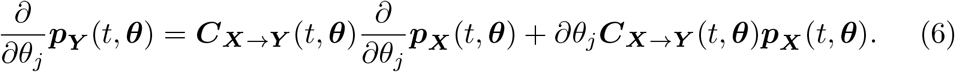

Then, the Fisher Information Matrix (FIM) ***F*** _***Y*** (*t*)_(***θ***) of the noisy measurements ***Y*** (*t*) at time *t* is computed by

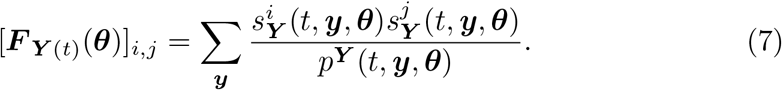

Details of the numerical approximation can be found in SI Section S3.1. In this formulation, we only need to solve the (usually expensive) sensitivity equations derived from the CME once, then apply relatively quick linear algebra operations to find the FIM corresponding to any new PDO for specific microscope, fixation protocol, or probe designs.

To convert the FIM between parameters defined in linear space to the same parameters in logarithmic space, we apply the transformation:

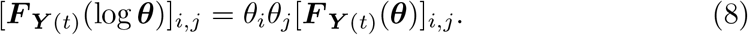

#### 2.1.5 Parameter estimation and uncertainty quantification

Models are fit to experimental data in Matlab, (R2021b, [35]) using an iterated combination of the builtin `fminsearch` algorithm (an implementation of the Neldar-Mead simplex method, [44]) to get close to the MLE (i.e., to maximize the likelihood in Eq. 5) followed by a customized version of the Metropolis Hastings (MH) sampling routine, `mhsample` ([14]). All parameter searches were conducted in logarithmic space, and model priors were defined as lognormal with log-means and standard deviations as described in the main text. For the MH sampling proposal function, we used a (symmetric) multivariate Gaussian distribution centered at the current parameter set and with a covariance matrix proportional to the inverse of the Fisher Information matrix (calculated at the MLE parameter set) and scaled by a factor of 0.8, which achieves an approximately 20–50% proposal acceptance rate for all combinations of PDOs and data sets. MH chains were run for 20,000 samples. Convergence was checked by computing the autocorrelation function and verifying that the effective sample size was at least 1000 for every parameter in every MH chain. All data, model construction, FSP analysis, FIM calculation, parameter estimation, and visualization tasks were performed using the Stochastic System Identification Toolkit (SSIT) available at: https://github.com/MunskyGroup/SSIT using manuscript specific scripts provided in folder `CommandLine/Vo et al 2023`.

### 2.2 Single-cell Labeling and Imaging

#### 2.2.1 Cell Culture

The experiments presented here were performed on Hela Flp-in H9 cells (H-128). The generation of the H-128 cell line has previously been discussed ([84]). Briefly, Tat expression regulates the MS2X128 cassette-tagged HIV-1 reporter gene in H-128 cells. The HIV-1 reporter consists of the 5′ and 3′ long terminal repeats, the polyA sites, the viral promoter, the SD1 and SA7 splice donors, and the Rev-responsive element. Additionally, the MS2 coating protein conjugated with a green fluorescent protein (MCP-GFP), which binds to MS2 stem loops when transcribed, is expressed persistently by H-128 cells. Cells were cultured in Dulbecco’s modified Eagle medium (DMEM, Thermo Fisher Scientific, 11960-044) supplemented with 10% fetal bovine serum (FBS, Atlas Biologicals, F-0050-A), 10 U/mL penicillin/streptomycin (P/S, Thermo Fisher Scientific, 15140122), 1 mM L-glutamine (L-glut, Thermo Fisher Scientific, 25030081), and 150 *μ*g/mL Hygromycin (Gold Biotechnology, H-270-1) in a humidified incubator at 37°C with 5% CO_2_.

#### 2.2.2 smiFISH and Microscopy

Single-molecule inexpensive fluorescence *in situ* hybridization (smiFISH) was performed following a protocol previously described ([88, 32]). This technique is known as inexpensive, because the primary probes consist of a region binding the transcript of interest plus a common sequence established as FLAP, which is bound by a complementary FLAP sequence conjugated with a fluorescent dye following a short PCR-cyle (the primary and FLAP-Y-Cy5 probes used in this study were purchased from IDT, see Table 1). To perform smiFISH, H-128 cells were plated on 18 mm cover glasses within a 12-well plate (*∼* 10^5^ cells/well), 24 h before the experiment. Some samples were exposed to 5 *μ*M triptolide (Sigma-Aldrich, 645900) for different incubation periods. Immediately after these drug treatment periods, samples were washed out twice with RNAse free 1XPBS, and fixed in 4% PFA at RT for 10 min, followed by 70% ethanol permeabilization at 4°C for least 1 h. After washing each sample with 150 *μ*L of wash A buffer (Biosearch Technologies, SMF-WA1-60) for 5 min, each cover glass was set on a droplet (cells facing down) consisting of 45 *μ*L hybridization buffer (Biosearch Technologies, SMF-HB1-10) and 1 *μ*L of the duplex smiFISH probes (MS2-transcript-binding probe mix + FLAP-Y-binding region annealed to FLAP-Y-Cy5) in a humidified chamber at 37°C overnight. The following day, samples were placed in a fresh 12-well plate, and the cells (facing up) were incubated twice in wash A buffer at 37°C for 30 min, first alone, and then containing DAPI. Finally, cells were incubated with wash B buffer (Biosearch Technologies, SMF-WB1-20) at RT for 5 min, and then mounted on a 15 *μ*L drop of Vectashield mounting medium (Vector Laboratories, H-1000-10), and sealed with transparent nail polish.

**Table 1:**
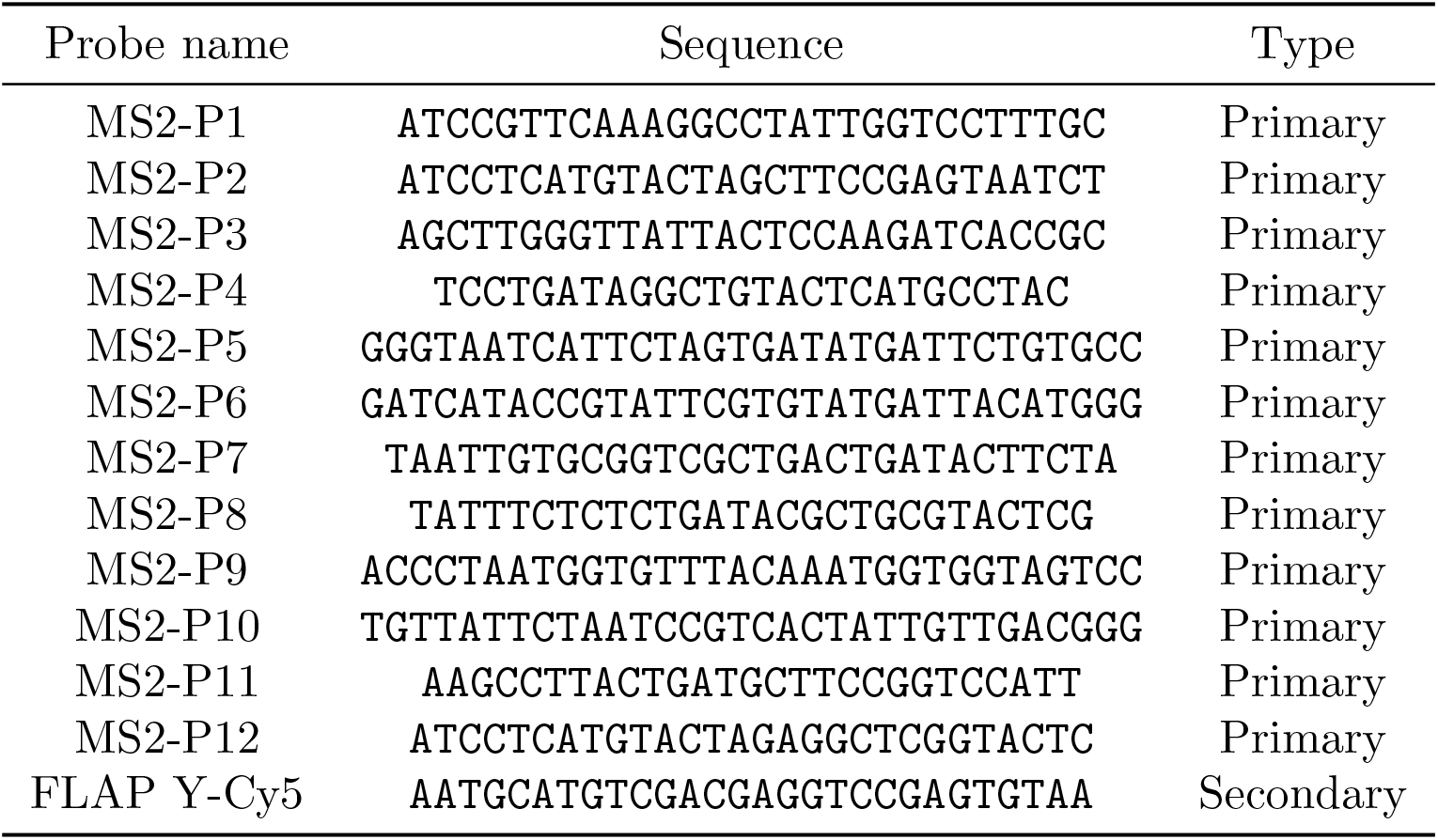
smiFISH probe sequences. Each primary probe has an added common FLAP-Y binding sequence (TTACACTCGGACCTCGTCGACATGCATT)

Fluorescent images were acquired with an Olympus IX81 inverted spinning disk confocal (CSU22 head with quad dichroic and additional emission filter wheel to eliminate spectral crossover) microscope with 60x/1.42 NA oil immersion objective. Confocal z-stacks (0.5 *μ*m step-size, 27 stacks in each channel) were collected. Each field of view was imaged using four high-power diode lasers with rapid (microsecond) switcher (405 nm for DAPI, 488 nm for MS2-MCP-GFP reporter, 561 nm for cytosol marker, and 647 nm for smiFISH MS2-Cy5, exposure time of 100 ms for all except smiFISH channel that was 300 ms) when samples had cytosol marker or three lasers for samples without the marker. The system has differential interference contrast (DIC) optics, built-in correction for spherical aberration for all objectives, and a wide-field Xenon light source. We used an EMCCD camera (iXon Ultra 888, Andor) integrated for image capture using Slide book software (generating 60× images with 160 nm/pixel). The imaging size was set to 624 × 928 pixels^2^.

#### 2.2.3 smiFISH image processing

In this study, we employed Python to implement an image processing pipeline comprising three steps: cell segmentation, spot detection, and data management (as previously described in [72]). Nuclear segmentation on the DAPI channel (405nm) was carried out using Cellpose ([82]) with a 70-pixel diameter as an input parameter. Spot quantification for both the MS2-MCP-GFP reporter channel (488 nm) and the smiFISH MS2-Cy5 channel (647 nm) was performed independently, using BIG-FISH software ([34]). The spot quantification procedure employed a voxel-XY of 160 nm, a voxel-Z of 500 nm, and spot radius dimensions of 160 nm and 350 nm in the XY and Z planes, respectively. This process included the following steps: i) the original image was filtered using a Laplacian of Gaussian filter to improve spot detection. ii) Local maxima in the filtered image were then identified. iii) To distinguish between genuine spots and background noise, we implemented both an automated thresholding strategy and a manual approach involving the use of multiple threshold values between 400 and 550. The intensity thresholds (550 for MS2-MCP-GFP and 400 for smiFISH MS2-Cy5) that resulted in the highest number of co-detected spots in both channels were chosen. Additionally, we quantified the average nuclear intensity for each color channel by computing the mean intensity of all pixels within the segmented nuclear region. Data management involved organizing all quantification data into a single dataset, including information on the specific image and cell in which the spots were detected. All image processing codes are available from https://github.com/MunskyGroup/FISH_Processing.

## 3 Results

Many models have been used to capture and predict observations of single-cell heterogeneity in gene expression ([58, 80, 15, 51, 59, 79, 75]). When selecting an experimental assay to parameterize such models, one is faced with several choices, each with its own characteristic measurement errors ([67]). Here, we start by introducing several mathematical forms for probabilistic distortion operators (PDOs) that can quantify these measurement errors. We then use a model and simulated data to show how different measurement errors can affect model identification, and we show how this can be corrected through consideration of the PDO in the estimation process. Next, we show how models and PDOs can be used in the framework of Fisher Information in iterative design of single-cell experiments for efficient identification of predictive models. Finally, we illustrate the practical use of the PDO, model inference, and FIM based experiment design on the experimental investigation of bursting gene expression from a reporter gene controlled by an HIV-1 promoter.

### 3.1 Distorted single-cell measurements sample a probability distribution that is the image of their true molecular count distribution through a linear operator

Most parameters needed to define single-cell signaling or gene expression models cannot be measured directly or calculated from first principles. Instead, these must be statistically inferred from datasets collected using single-molecule, single-cell experimental methods such as smFISH ([64, 29]), flow cytometry ([48, 50]), or live-cell imaging ([83, 24, 21]). In this work, we focus on the former two experimental approaches in which collected data consists of independent single-cell measurements taken at different times. Here, we consider measurement distortion effects corresponding to probe binding inefficiency and spot detection for smiFISH experiments, and in the supplemental text, we extend this to consider effects of reporter fluorescence intensity variability in flow cytometry experiments (Section S3.1), data binning (Section S3.2), effects of competition with non-specific probe targets (Section S3.3), and effects of segmentation errors (Section S3.4).

We first consider five formulations to define the measurement distortion matrix (cf. eq. (4) in Methods, and illustrated in Fig 2), corresponding to scenarios in which experimental errors arise from either inefficient mRNA detection, additive false positives, or combinations. Specifically,

**Figure 2:**
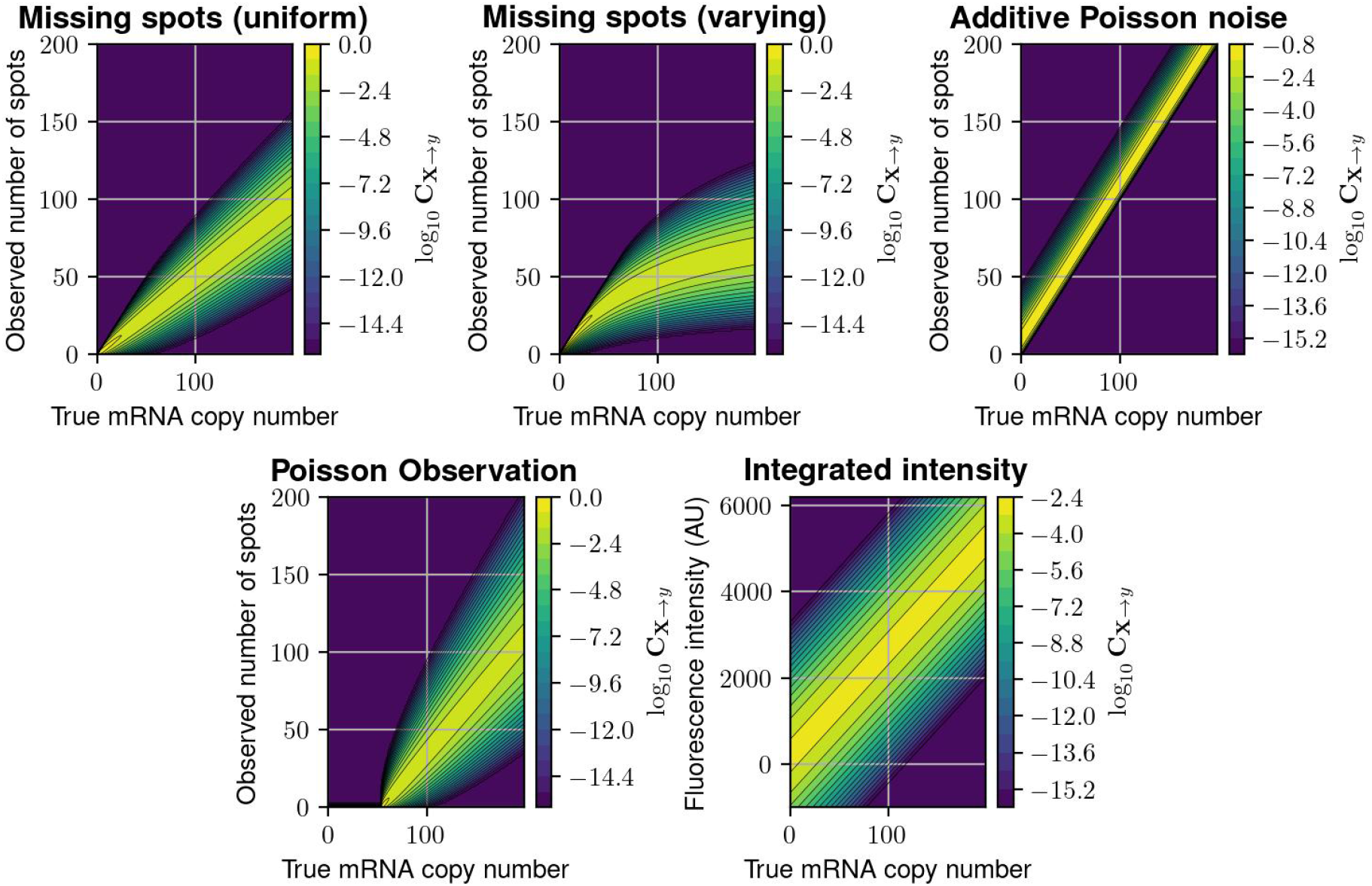
Probabilistic distortion operators. **(A)** The “Missing Spots” (MS) distortion model, where spots can randomly go missing. **(B)** The “Missing Spots with Variable Rate” (MSVR) distortion, where the probability of missing a spot increases with spot density. **(C)** The “Poisson Noise” (PN) model, where false positive spots are added to the counted number of spots. **(D)** A “Poisson Observation” (PO) model, where the detected spots follow a Poisson distribution with mean proportional to the true number. **(E)** The “Integrated Intensity” (II) model, where only a perturbed version of the total fluorescence is recorded per cell. In each heat map, the color at point (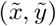) is the conditional probability mass/density of the measurement *y* having value 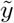 given that the true copy number *x* has value 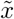.

1. The first model supposes that 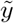 is obtained from a “lossy” spot-counting process applied on images taken in an smFISH experiment. We model *y*|***x*** = *j* with a binomial distribution *B*(*p*_detect_, *j*), where each spot has a chance *p*_miss_ := 1 – *p*_detect_ of being ignored by the counting algorithm, resulting in underestimation of the true mRNA copy number. In the context of optical microscopy, such a distortion might result from quantifying spots at a single plane, where *p*_detect_ might represent the fraction of the imaged section compared to the full volume of the cell, but similar error models have also been proposed in the context of single-cell RNA sequencing ([45, 6]). We call this distortion “Missing Spots”. Its PDO, **C**_MS_ is illustrated in Fig 2A for *p*_miss_ := 0.5 and can be defined:

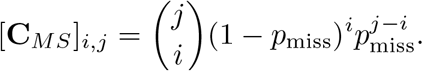
2. The second model is a simple variation of the first model in which *p*_detect_(*j*) varies with the number of mRNA molecules *j*. For example, the specific formulation, *p*_detect_ := 1.0/(1.0 + 0.01*j*) implies that spot detection rate degrades as the number of mRNA molecules in the cell increases, which may correspond to the effect of co-localization and/or image pixelation which could cause the under-counting of overlapping spots. We call this model “Missing Spots with Varying Rate”, and **C**_MSVR_ is illustrated in Fig 2B and can be expressed:

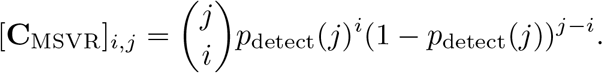
3. The third model assumes that 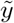 is the output of a spot-detection process contaminated by false positives, e.g., due to background fluorescence noise in the image that can appear to be spots. We model these false positives by additive Poisson noise, making 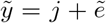 where 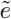 ∼Poisson(*λ*). We call this model “Poison Noise,” and **C**_PN_, which is illustrated in Fig 2C for *λ* = 10 and can be expressed:

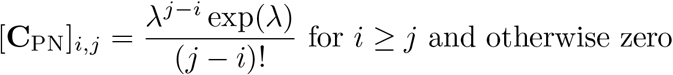
4. The fourth model is a simple extension of the third model in which the number of detected spots given a true number, *j*, is a Poisson distribution with a varying mean *λ*(*j*). Specifically, *λ*(*j*) := max (0, *λ*_0_ + *λ*_1_*j*), where *λ*_0_ is Poisson noise, and *λ*_1_ is the Poisson detection rate. We call this model “Poisson Observation”, and **C**_PO_ is illustrated in Fig 2D and expressed as:

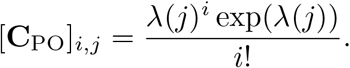
5. The fifth model concerns fluorescent intensity integration measurements (such as those used in flow cytometry). We use the model proposed in ([58]), in which 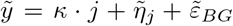, where 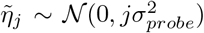 models fluorescent heterogeneity and 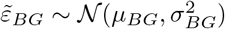 is the background noise. The PDO is a hybrid matrix with discrete columns (i.e., its domain are probability vectors over discrete CME states) and continuous rows (its range consists of continuous probability density functions over the range of fluorescence intensities). This PDO, which we label “Integrated Intensity” and is illustrated in Fig 2E for *μ*_*BG*_ = 200, *σ*_*BG*_ = 400, 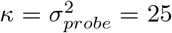, can be expressed:

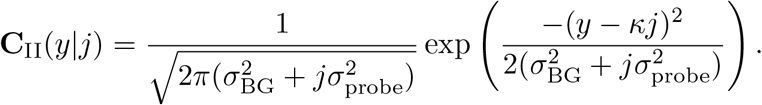

For intensity measurement with finite resolution, **C**_II_ can be defined over discrete bins (e.g., (*y*_0_, *y*_1_], (*y*_1_, *y*_2_], …) by integrating as follows:

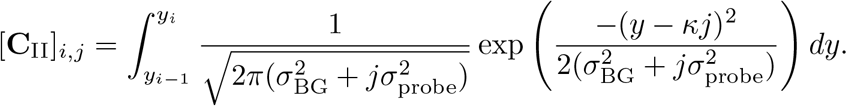

We stress that these PDOs are provided as just a few of many possible distortions that could be modeled using the proposed framework. Other, more complex distortion operators are discussed in Supplemental Text S3, including one where a secondary CME is employed to model the uncertain chemical kinetics of the measurement process (Section S3.3) and another to describe cell merging due to image segmentation errors (Section S3.4).

#### 3.1.1 Measurement noise introduces bias and uncertainty into model identification

To illustrate the impact of measurement distortion on parameter estimation and our use of the PDO formalism to mitigate these effects, we begin with an analysis of the random telegraph model (Fig 3A), one of the simplest, yet most commonly utilized models of bursty gene expression ([61, 64, 83, 73, 74, 45]). In this model, a gene is either in the inactive or active state, with transition between states occurring randomly with average rates *k*_ON_ (to activate the gene) and *k*_OFF_ (to deactivate the gene). When active, the gene can be transcribed with an average rate *k*_*r*_ to produce mRNA molecules, each of which degrades with an average rate *γ*.

**Figure 3:**
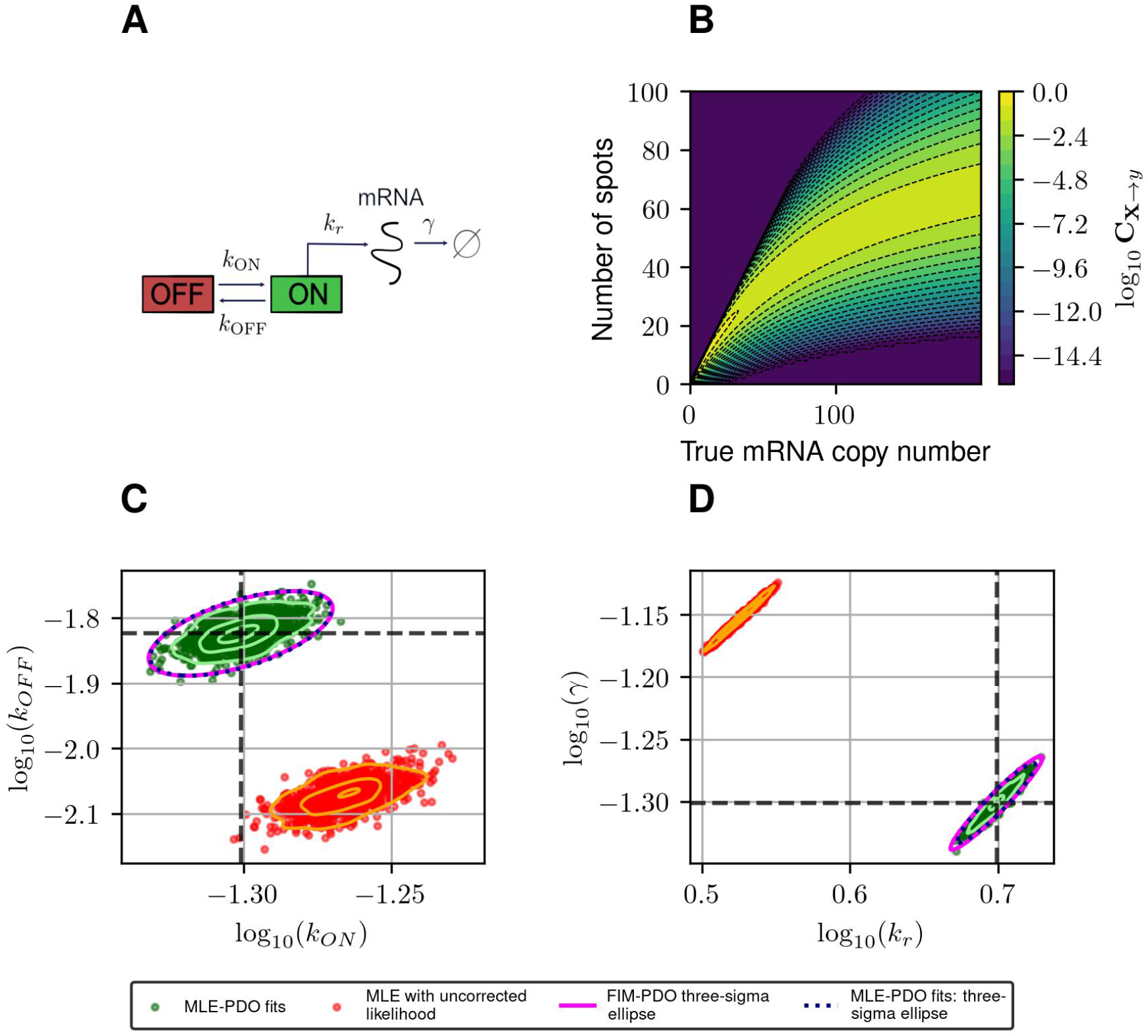
Estimating and correcting for how measurement distortion affects model identification. **(A)**: Schematic of the random telegraph gene expression model. Parameter values: gene activation rate *k*_*ON*_ = 0.015 events/minute, gene deactivation rate *k*_*OFF*_ = 0.05 events/minute, transcription rate *k*_*r*_ = 5 molecules/minute, degradation rate *γ* = 0.05 molecules/minute. **(B)**: A submatrix of the PDO for missing spots with varying rates, restricted to the domain {0, …, 200}×{0, …, 100}. **(C) and (D)**: Maximum likelihood fits to validate FIM-based uncertainty quantification for observed mRNA distributions under distortion model shown in (B). We simulated 1, 000 datasets and perform maximum likelihood fits to these datasets using either a likelihood function that ignores measurement noise (red, labeled ‘MLE with uncorrected likelihood’), or one whose measurement noise is corrected by incorporating the PDO (dark green, labeled ‘MLE-PDO fits’). The estimated density contours (delineating 10, 50, and 90 -percentile regions) of the fits are superimposed in light shades. Also displayed are the three-sigma confidence ellipses computed by the FIM-PDO approach or from sample covariance matrix of the corrected MLE fits. Panel (C) shows the results in log_10_(*k*_*OFF*_) – log_10_(*k*_*ON*_) plane, while panel (D) shows them in the log10(*k*_*r*_) log_10_ (*γ*) plane. The intersection of the thick horizontaland vertical lines marks the location of the true data-generating parameters. See Table 2 for a quantitative comparison between the uncorrected MLE and MLE-PDO.

To demonstrate how measurement distortions affect parameter identification, and why explicit measurement error modeling is necessary, we use the bursting gene expression model to simulate mRNA expression data where each cell is distorted by the MSVR effect above (PDO reproduced in Fig 3B). Each data set consists of five batches of 1,000 independent single-cell measurements that are collected at five equally-spaced time points *j*Δ*t, j* = 1, 2, 3, 4, 5 with Δ*t* = 30 minutes. We considered two methods to fit the telegraph model to these data based on the Maximum Likelihood Estimator (MLE): one in which the likelihood function ignores measurement noise, and one where measurement noise modeled by the PDO is incorporated into the likelihood function (see Supplemental text S2 for their formulations). If one fails to account for measurement uncertainty, these fits produce strongly biased estimates for the RNA production and degradation rates as seen in Fig 3C,D (red). Because this bias is inherent to the measurement technique, it cannot be corrected simply by averaging over more experiments. On the other hand, using the distortion correction method, Fig 3C,D (dark green) shows that the inaccurate spot counting procedure can be corrected by explicitly accounting for measurement uncertainty in the modeling phase. Quantitatively (see Table 2), the MLE fits that incorporate noise modeling have lower bias (in terms of relative root-mean-squared errors) for all four parameters compared to the uncorrected fits.

**Table 2:**
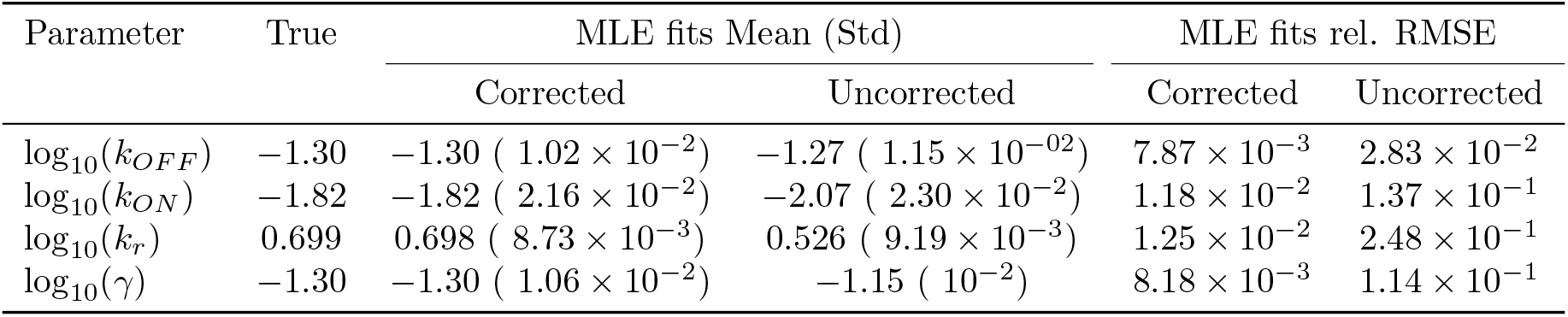
Performance of the maximum likelihood estimator (MLE) for estimating bursting transcription kinetic parameters. The third and fourth columns compare the mean and standard deviation of fits with and without PDO correction (labeled Corrected and Uncorrected, respectively). The final two columns compare the relative root-mean-squared errors (RMSEs) of these fits. For a quantity of interest *q* and its *n* estimated values 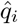, *i* = 1, …, *n*, we define the relative RMSE as 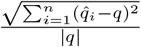.

| Parameter | True | MLE fits Mean (Std) |  | MLE fits rel. RMSE |  |
| --- | --- | --- | --- | --- | --- |
|  |  | Corrected | Uncorrected | Corrected | Uncorrected |
| $\log_{10}(k_{OFF})$ | -1.30 | -1.30 ( $1.02 \times 10^{-2}$ ) | -1.27 ( $1.15 \times 10^{-02}$ ) | $7.87 \times 10^{-3}$ | $2.83 \times 10^{-2}$ |
| $\log_{10}(k_{ON})$ | -1.82 | -1.82 ( $2.16 \times 10^{-2}$ ) | -2.07 ( $2.30 \times 10^{-2}$ ) | $1.18 \times 10^{-2}$ | $1.37 \times 10^{-1}$ |
| $\log_{10}(k_r)$ | 0.699 | 0.698 ( $8.73 \times 10^{-3}$ ) | 0.526 ( $9.19 \times 10^{-3}$ ) | $1.25 \times 10^{-2}$ | $2.48 \times 10^{-1}$ |
| $\log_{10}(\gamma)$ | -1.30 | -1.30 ( $1.06 \times 10^{-2}$ ) | -1.15 ( $10^{-2}$ ) | $8.18 \times 10^{-3}$ | $1.14 \times 10^{-1}$ |

Figures 3C,D also demonstrate that the Fisher Information Matrix computed for the noisy measurement (see eq. (7)) produces a close approximation to the covariance matrix of the MLE (compare magenta and black ellipses). This suggests that the FIM can provide a relatively inexpensive means to quantify the magnitude and direction of parameter uncertainty, a fact that will be helpful in designing experiments that can reduce this uncertainty, as we will explore next.

#### 3.1.2 Fisher Information Matrix Analysis reveals how optimal experiment design can change in response to different measurement distortions

Having demonstrated the close proximity between our FIM computation and MLE uncertainty for simulated analyses of the bursting gene expression model, we next ask whether the sampling period Δ*t* could be tuned to increase information but using the same number of measurements. Recall that our simulated experimental set-up is such that measurements could be placed at five uniformly-spaced time points *t*_*j*_ = *j*Δ*t, j* = 1, 2, 3, 4, 5, with the sampling period Δ*t* in minutes, and that at each time point we collect an equal number *n* of single-cell measurements, chosen as *n* := 1000. We find the optimal sampling period Δ*t* for each measurement distortion (MS, MSVR, PN, PO, and II), and compare the most informative design that can be achieved for each class. Here, we define ‘optimal’ in terms of the determinant of the Fisher Information Matrix, the so-called D-optimality criterion, whose inverse estimates the volume of the parameter uncertainty ellipsoid for maximum likelihood estimation ([2]). Figures 4A,B shows the information volume to the five kinds of noisy measurement described above (Fig 2), in addition to the ideal noise-free smiFISH, at different sampling rates. We observe that every probabilistic distortion to the measurement decreases the information volume (but to different extents), and that each measurement method results in a different optimal sampling rate. In Fig 4C-D, we plot the three-sigma (i.e., 99.7%) confidence ellipsoids of the asymptotic distribution of MLEs projected on the log_10_(*k*_*ON*_) − log_10_(*k*_*OFF*_) plane and the log_10_(*k*_*r*_) − log_10_(*γ*) plane.

**Figure 4:**
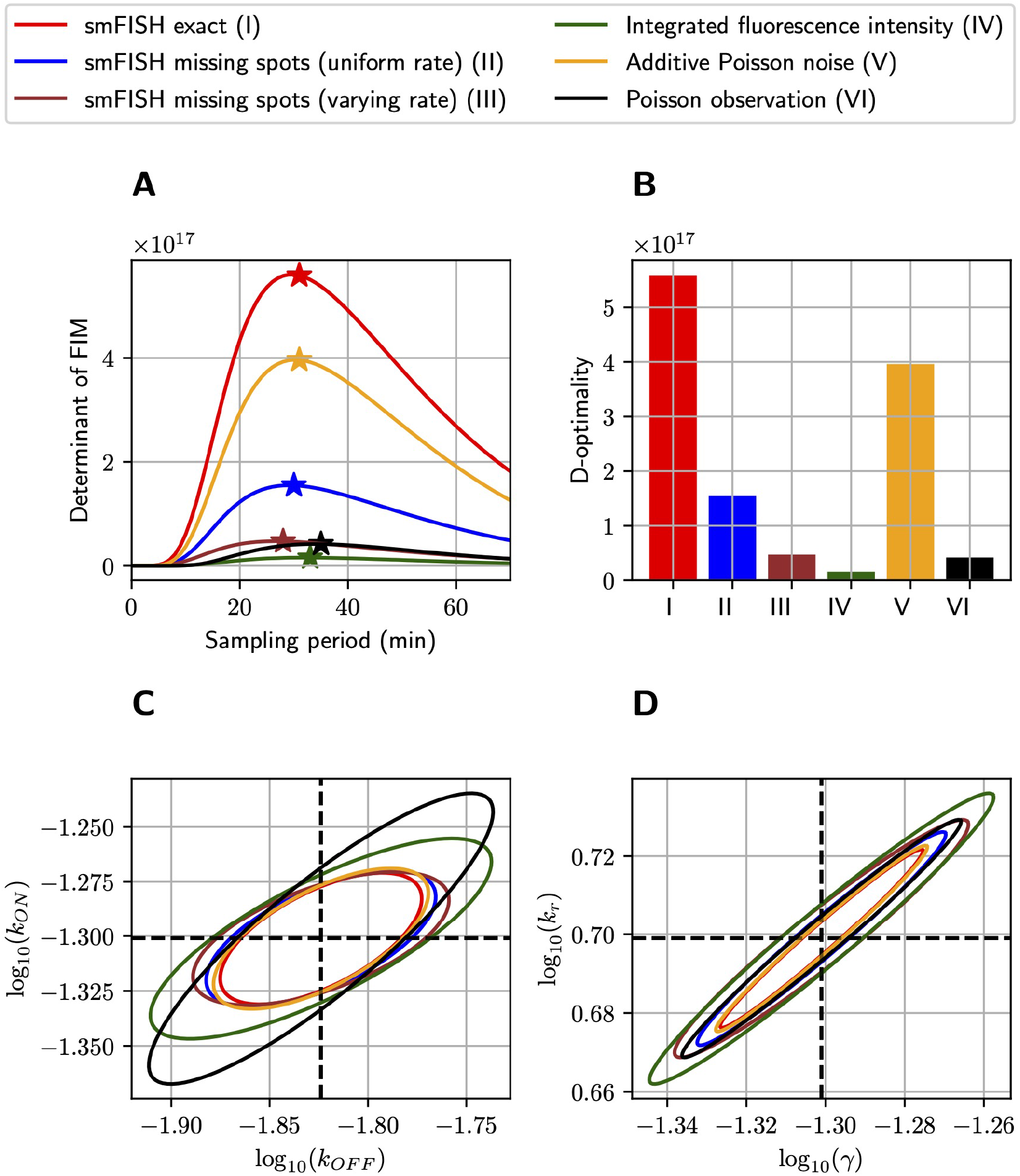
Optimizing experiment sampling rate under different measurement distortion effects. **(A)**: Comparison of D-optimality criteria in single-cell experiments with different types of measurement noise and at different sampling intervals (Δ*t*). In this settings, independent measurements are collected at five equally-spaced time points *k*Δ*t, k* = 1, 2, 3, 4, 5 with 1000 measurements placed at each time-point. The ⋆ symbol marks the optimal point of each curve and the bar charts in **(B)** visualizes the relative differences in D-optimal achievable by the measurement methods at their respective optimal sampling rates. **(C) and (D)**: The three-sigma confidence ellipses projected onto the log_10_(*k*_*ON*_)–log_10_ (*k*_*OFF*_) and log_10_ (*k*_*r*_) –log_10_ (*γ*) planes. These ellipses are computed by inverting the FIMs of different measurement noise conditions at their optimal sampling rates. All analyses use the model and parameters from Fig 3.

### 3.2 FIM and PDO Analysis of Experimental Measurements for HIV-1 Promoter Bursting Kinetics

To provide a concrete example for the use of the FIM and PDO in practice, we performed single-cell measurements to quantify the relative measurement distortion between different single-mRNA labeling strategies in HeLa (H-128) cells (see Methods). We expressed a transcription reporter gene with 128 repeats of the MS2 hairpin, and we simultaneously used both MCP-GFP (green labels in Fig 5A) and smiFISH MS2-Cy5 (magenta labels in Fig 5A) to target the MS2 repeats. In each cell, both approaches detect similar patterns for the number and spatial locations of mRNA within the nuclei (Fig 5B-G). For our particular choice of image processing algorithm ([72, 82, 34]) and intensity threshold for spot detection (see Methods) and a two-pixel (x,y,z) Euclidean distance threshold for co-localization detection, after analyzing 135 cells in steady-state conditions, we found that 46.9% of mRNA spots (15,258 out of 32,526 total) were detected in both channels (e.g., spots denoted by white triangles in Fig 5A). However, many spots (21.3%, 6,913 spots) are detected only using smiFISH (e.g., those denoted with magenta triangles) and 31.8% (10,355 spots) are only detected using the MS2-MCP labels (e.g., green triangles). Due to differences in label chemistry, background fluorescence, and image analysis errors, the quantified distribution of mRNA expression depends heavily upon which assay is utilized (compare Figs 5C-E to 5F-H). For example, we observed that analyses of cells that have fewer than 10 spots in the smiFISH channel frequently result in large numbers of spurious spots when the same cells are analyzed in the MCP-GFP channel (black markers on left limit of Fig 5C and high density at zero in Figs 5D,E). These differences in measurement quantification support the need for consideration of measurement distortions in subsequent analysis.

**Figure 5:**
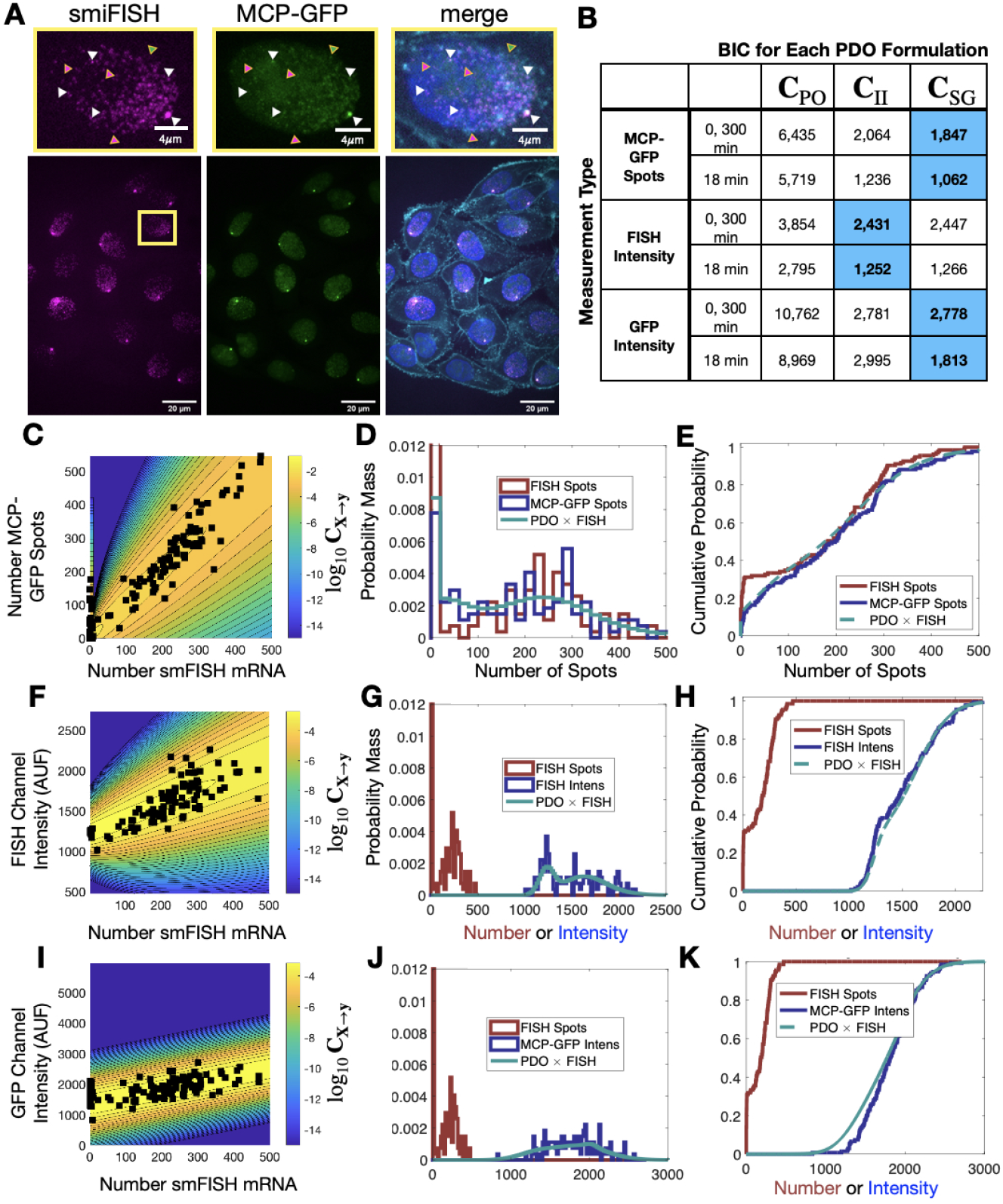
Effect of labeling strategy on single-cell mRNA expression quantification. **A**) Example image of a single-cell population expressing a reporter gene controlled by an HIV-1 promoter and containing 128*X*MS2 stem-loop cassette, in which the mRNA was simultaneously measured using MCP-GFP labeling (green) and with smiFISH probes against MS2-Cy5 (magenta). A higher resolution image of the indicated single nucleus is shown at the top; merged image on the right includes DAPI and MemBrite cell surface stain 543/560 to denote the cell nucleus and plasma membrane, respectively. Triangles denote example mRNA that are detected in both channels (white, 34.8%), only in the MCP-GFP channel (green, 37.5%) and only in smiFISH (MS2-Cy5) channel (magenta, 27.7%). **B**) BIC of different combinations of PDO (columns) and measurement type (rows) given an assumed ‘true’ measurement of smiFISH mRNA. In all cases, PDO parameters are chosen to maximize likelihood for *t* = 0 and 300 min data, and *t* =18 min is predicted without changing parameters. Blue shading denotes PDO selection is identical if based on BIC for (0,300) min data or prediction of 18 min data. **C**) Scatter plot of the spot count using MCP-GFP versus using smiFISH for data collected at *t* =0 min (black squares). Shading and contour lines denote the levels of the PDO (log_10_ ***C***) determined empirically from the data. **D**,**E**) Empirical probability mass (D, bin size = 20) and cumulative distributions (E, no binning) for smiFISH spot count (red) and MCP-GFP spot count (blue). Predicted MCP-GFP spot distributions using the smiFISH spot measurements and the estimated PDO are shown in green. **F-H**) Same as (C-E) but for the total integrated intensity measurement of the smiFISH channel. **I-K**) Same as (C-E) but for the total integrated intensity measurement of the GFP channel.

#### 3.2.1 PDO measurement noise parameters can be calibrated using single-cell experiments with multiple measurement modalities

To demonstrate the parameterization and selection of a PDO for these data, we measured expression using smiFISH or MCP-GFP spot counts as well as with total integrated fluorescence intensity in the FISH or GFP channels. These measurements were collected for 135 cells at t=0 and 62 cells at t=300 min after transcription deactivation by 5*μ*M Trp. We then defined the detected smiFISH spots as the “true” measurements {*x*_1_, …, 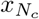 } and considered the MCP-GFP spot counts, FISH intensity, and GFP intensity values as three different groups of distorted observations {*y*_1_, …, 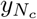 }. We then assumed three possible PDO formulations for each measurement modality, including the simple “Poisson Observation” (PO) and “Integrated Intensity” (II) PDOs from above, as well as an extended “Spurious Gaussian” (SG) PDO that is formulated starting with the II model but then extended to allow for a random fraction (*f*) of cells with a true spot count of 10 or fewer cells to be miscounted as a random observation drawn from a Gaussian (*i ∼ N* (*μ, σ*^2^)) distribution. For all PDOs, we assumed that distorted quantities would be rounded to their nearest non-negative integer value (e.g., negative values would be rounded up to zero). For a each PDO and its corresponding parameter set **Λ**, we calculate the corresponding PDO, **C**(**Λ**), and the log-likelihood to observe {*y*_*i*_} given {*x*_*i*_ ≥ is computed as

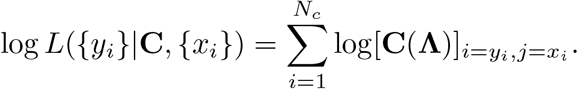

We then independently maximize this likelihood function for each combination of the three different distorted data sets (MCP-GFP spots, FISH intensity, GFP intensity) and for the three PDO formulations (PO, II, SG). For each data set, we finally select the PDO formulation that results in lowest Bayesian Information Criteria (BIC ≡ *k* log(*N*_*c*_) − 2 log *L*, where *k* is the number of parameters in the PDO) for the t=0 and t=300 min data (Fig 6B). We also verified in all cases that the PDO selection also maximized the likelihood of the predictions for held-out data at t=18 min after Tpt treatment (Fig 6B). Upon fitting and selection based on either BIC or cross-validation, we found that MCP-GFP spot count data and GFP intensity distortions were best represented by the “Spurious Gaussian” PDO (**C**_SG_), while the FISH intensity measurements were best represented by the “Integrated Intensity” PDO (**C**_II_).

**Figure 6:**
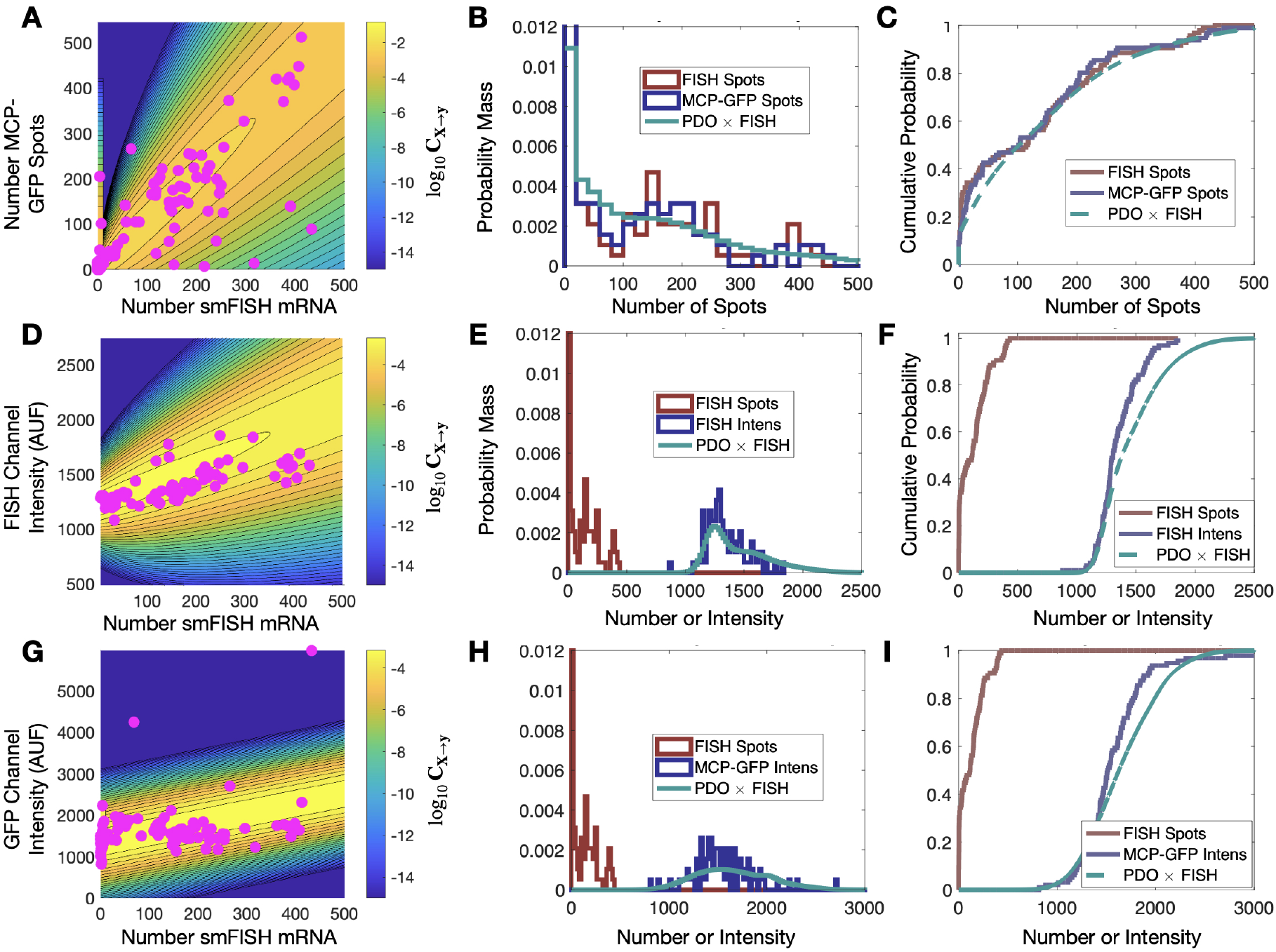
Validation of PDO on held out data. **A**) Scatter plot of smiFISH spot counts and MCP-GFP spot counts for data collected at *t* =18 min and contour of PDO determined from data at *t* = (0, 300) min. **B**,**C**) PDF (bin size = 20) and CDF for smiFISH mRNA detection (red) or with MCP-GFP (blue) and MCP-GFP-based PDO-prediction of total mRNA (green). **D-F**) Same format as (B-D) but for measurements of the total smiFISH fluorescence intensity per cell.**G-I**) Same format as (B-D) but for measurements of the total GFP fluorescence intensity per cell.

Figures 5C,F show the contours of the corresponding PDOs for the MCP-GFP spot and FISH intensity data, respectively, and Figs 5D,E,G,H show the PDF and CDF for the “true” smiFISH mRNA count data data at t=0 (blue lines) compared to the distorted data (red lines). From the figures, we find the distortion model does a good job to calibrate between the total mRNA and MCP-GFP-or smiFISH-detected spot counts (compare blue and green lines in Figs 5D,E,G,H and the BIC values for (0,300) min in Fig 5B). We next verified that the PDO remains constant by showing that the same models and same parameters also accurately reproduce the difference in total mRNA and MCP-GFP or smiFISH measurements at a held out time of 18 min after application of 5 *μ*M Trp (see Fig 6, and BIC values for 18 min in Fig 5B).

#### 3.2.2 The PDO allows estimation of predictive bursting gene expression model parameters from distorted data

We next asked if using the estimated PDO while fitting the MCP-GFP spot data or the total FISH or GFP intensity data would enable the identification of appropriate model parameters to predict the smiFISH mRNA counts. Based on previous observations ([21]), we proposed a 3-state bursting gene expression model (Fig 7A) where each of two alleles can occupy one of three states: (S1=OFF) where no transcription can occur; (S2=Poised) where the promoter is ready to begin transcription; and (S3=Active) where transcripts are produced in rapid bursts of mRNA expression. The model contains six parameters: *k*_ON_ and *k*_OFF_ are the promoter transition rates between the OFF and Poised states; *ω* and *k*_Ex_ are the burst frequency and burst exit rates; and *k*_*r*_ and *γ* are the transcription and mRNA degradation rates, respectively. Based on previous observation ([21]) that triptolide (Trp) represses transcription after an average of 5-10 min needed for diffusion of Trp to the promoter and completion of nascent mRNA elongation and processing, we model the Trp response as a complete inactivation of transcription (*ω* → 0) that occurs at t=5 min.

**Figure 7:**
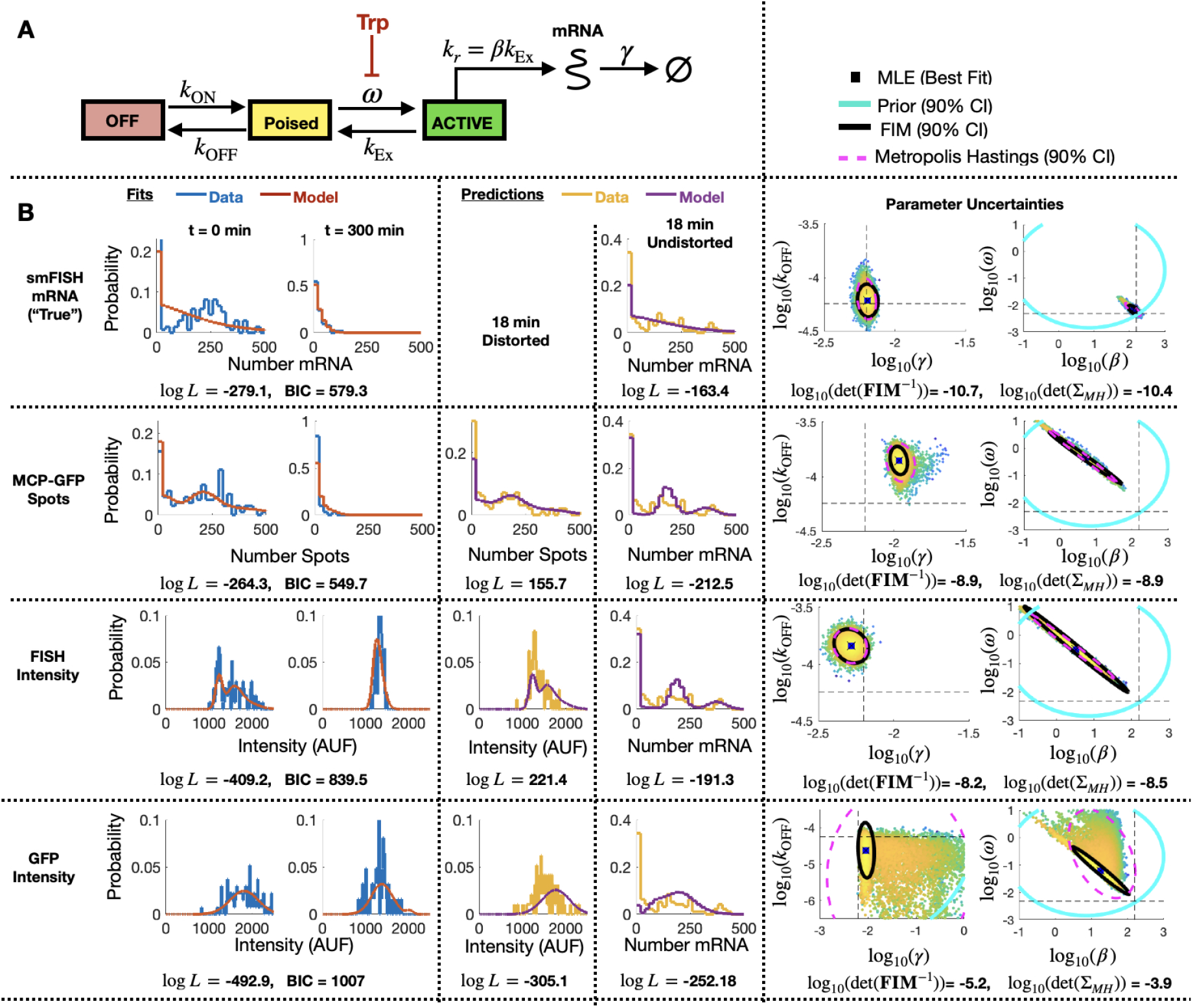
Identification of stochastic model for MS2X128 cassettetagged HIV-1 reporter gene. **(A)** Schematic of the 3-state bursting gene expression model. **(B)** Results for model fitting, prediction, and uncertainty quantification for measurements based smiFISH spots (top row), MCP-GFP spots (row 2), total FISH intensities (row 3) and GFP intensities (row 4). Left two columns show the measured and model-fitted probability mass vectors (PMV) at 0 and 300 min after 5*μ*M Tpt. Third column shows the model-predicted and measured PMV for the corresponding (distorted) measurement modality at 18 min after 5*μ*M Tpt. Fourth column shows the model prediction without distortion and measured PMV for the smFISH mRNA count at 18 min after 5*μ*M Tpt. All histograms use a bin size of 20. Log-likelihood values for all model-data comparisons (and BIC values for fitting cases, *k* = 4 parameters, N = 197 cells) are computed without binning and are shown below the corresponding histograms. Right two columns show joint parameter uncertainty for model estimation using data for 0 and 300 min after 5*μ*M Tpt. In each case, the 90% CI for prior is shown in cyan; Metropolis Hastings samples (N = 20,000) are shown in dots; 90% CI for posterior is shown in dashed magenta; and FIM-based estimate of 90% CI is shown in black. Horizontal and vertical dashed black lines denote the “true” parameters and are defined as the MLE when using fit to the smFISH counts and using all time points. Determinant of inverse FIM and covariance of MH samples is shown below each pair of uncertainty plots (both use log base 10).

To specify a prior guess for the model parameters, we considered published values from ([21]), where we used live-cell imaging of ON and Poised transcription sites to determine the burst frequency *ω* ≈ 0.2 min^−1^ and that *k*_*r*_ and *k*_Ex_ are too fast to be estimated independently, but are related by a burst size of *k*_*r*_/*k*_Ex_ = *β* ≈ 7.1 mRNA/burst. Also, based on observations in ([21]) that transcription sites remain in the OFF or Poised/Active states for long periods of 200 minutes or more, we assumed that *k*_ON_ and *k*_OFF_ would be too slow to estimate except under much longer experiments. We therefore estimated *k*_ON_ = *k*_OFF_ = 10^−4^ min^−1^, but we sought only to estimate the relative rate for *k*_OFF_. We estimated a typical mammalian mRNA half-life of 60 min yielding a rate *γ* = 5.8 × 10^−3^ min^−1^. With these baseline values as rough estimates, we assumed a log-normal prior distribution with a standard log deviation of one order of magnitude from the literature-based values for parameters {*ω, β, γ*} and two orders of magnitude for the more approximate value for *k*_OFF_}. These initial parameter guesses and prior uncertainties are summarized in Table 3(column 3).

**Table 3:** Parameter priors, MLE estimates, Uncertainties upon Initial Fit. Initial estimates after fitting to data at *t* =(0,300) min. MH results are from a chain of 20,000 samples. All parameter values and standard deviationsvalues are shown in log_10_.

| Parameter | Quantity | Prior | smiFISH | MCP-GFP Spots | FISH Intens | GFP Intens |
| --- | --- | --- | --- | --- | --- | --- |
| $k_{\text{OFF}} \text{ (s}^{-1}\text{)}$ | $\log_{10}(\lambda)$ | -4.0 | -4.215 | -3.852 | -3.840 | -4.619 |
| | $\sigma_{p/MH}$ | 2.0 | 0.091 | 0.083 | 0.069 | 1.160 |
| | $\sigma_{FIM}$ | 2.0 | 0.071 | 0.060 | 0.070 | 0.308 |
| $\omega \text{ (s}^{-1}\text{)}$ | $\log_{10}(\lambda)$ | -0.70 | -2.173 | -0.357 | -0.505 | -1.205 |
| | $\sigma_{p/MH}$ | 1.0 | 0.117 | 0.462 | 0.559 | 0.752 |
| | $\sigma_{FIM}$ | 1.0 | 0.077 | 0.356 | 0.475 | 0.335 |
| $\beta \text{ (s}^{-1}\text{)}$ | $\log_{10}(\lambda)$ | 0.85 | 2.110 | 0.710 | 0.524 | 1.262 |
| | $\sigma_{p/MH}$ | 1.0 | 0.104 | 0.451 | 0.556 | 0.428 |
| | $\sigma_{FIM}$ | 1.0 | 0.069 | 0.379 | 0.438 | 0.307 |
| $\gamma \text{ (s}^{-1}\text{)}$ | $\log_{10}(\lambda)$ | -2.2 | -2.191 | -1.965 | -2.283 | -2.036 |
| | $\sigma_{p/MH}$ | 1.0 | 0.031 | 0.043 | 0.053 | 0.695 |
| | $\sigma_{FIM}$ | 1.0 | 0.034 | 0.028 | 0.053 | 0.078 |

**Table 4:** Parameter priors, MLE estimates, Uncertainties upon Final Fit. Final estimates after fitting to data at *t* =(0,18,300) min. MH results are from a chain of 20,000 samples. All parameter values and standard deviations values are shown in log_10_.

| Parameter | Quantity | Prior | smiFISH | MCP-GFP Spots | FISH Intens | GFP Intens |
| --- | --- | --- | --- | --- | --- | --- |
| $k_{\text{OFF}} \text{ (s}^{-1}\text{)}$ | $\log_{10}(\lambda)$ | -4.0 | -4.245 | -3.734 | -3.711 | -4.761 |
| | $\sigma_{p/MH}$ | 2.0 | 0.066 | 0.050 | 0.063 | 0.962 |
| | $\sigma_{FIM}$ | 2.0 | 0.066 | 0.045 | 0.060 | 0.506 |
| $\omega \text{ (s}^{-1}\text{)}$ | $\log_{10}(\lambda)$ | -0.70 | -2.318 | -0.105 | -0.891 | -0.591 |
| | $\sigma_{p/MH}$ | 1.0 | 0.066 | 0.478 | 0.533 | 0.396 |
| | $\sigma_{FIM}$ | 1.0 | 0.071 | 0.368 | 0.373 | 0.247 |
| $\beta \text{ (s}^{-1}\text{)}$ | $\log_{10}(\lambda)$ | 0.85 | 2.194 | 0.453 | 0.834 | 1.467 |
| | $\sigma_{p/MH}$ | 1.0 | 0.059 | 0.476 | 0.522 | 0.317 |
| | $\sigma_{FIM}$ | 1.0 | 0.057 | 0.427 | 0.454 | 0.227 |
| $\gamma \text{ (s}^{-1}\text{)}$ | $\log_{10}(\lambda)$ | -2.2 | -2.200 | -1.980 | -2.322 | -1.186 |
| | $\sigma_{p/MH}$ | 1.0 | 0.035 | 0.034 | 0.061 | 0.101 |
| | $\sigma_{FIM}$ | 1.0 | 0.036 | 0.028 | 0.062 | 0.072 |

Having specified parameter priors, we then took single-cell data for 135 cells at steady state (t=0) and 62 cells at t=300 min after application of 5*μ*M Trp, and we applied the Metropolis-Hasting (MH) algorithm (20,000 samples) to estimate rates and uncertainties for the four free parameters. As above, we assumed that the total spot count analysis provided the “true” spot count, and we considered four estimation problems using either the “true” smiFISH mRNA counts, the MCP-GFP spot count data, the FISH intensity data, or the GFP intensity data, each using the empirically estimated distortion operators from before (e.g., Figs 5C,F,I for the MCP-GFP spots, smiFISH intensity, and GFP intensity data, respectively). Table 3 presents the MLE parameter values after this initial stage, and Fig 7B (left two columns) compare the resulting fits of the model to the data. Figure 7B (right columns) shows scatter plots of parameter uncertainties for the parameter identification using either the smiFISH data (top row) or with the three distortion measurements using the empirical distortion operator (bottom three rows). From Figs 7B and Table 3, one can see that all fits do a reasonable job to match their intended data, and when fit using the PDO parameters, model predictions based only on the smiFISH and MCP-GFP data match well to the total spot count analysis (Fig 7B, middle column).

#### 3.2.3 The PDO-corrected FIM accurately estimates parameter uncertainty after experimental analysis of the bursting gene expression model

We next used the maximum likelihood models when fit to the 0 and 300 min data to compute the FIM for the corrected MCP-GFP and smiFISH measurements. Because the parameters cover multiple orders of magnitude, we transform the FIM to consider the parameters in log10 space (Eq 8). To adjust the FIM to consider the case where there is a prior on the parameters, we add the inverse of the prior covariance (in logspace) to the calculated FIM: 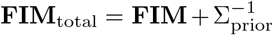. Given that we only collected a single experimental data set for each time point, it is not possible to directly compare the FIM to the spread of MLE estimates like we could for simulated data. However, in Fig 7B (right columns) for the smiFISH mRNA count and PDO-corrected MCP-GFP spots/intensity or smiFISH intensity data, we can compare the uncertainty predicted by the FIM analysis to the posterior uncertainty of our parameters given our data. This comparison shows that in most cases the FIM does an excellent job to estimate the direction and magnitude of parameter uncertainties (compare overlapping black and magenta ellipses in Figs 7B and see Table 3 for direct comparison of estimated standard deviations). However, it is important to note an exception where the FIM prediction does not match the MH analysis for the largest distortion (GFP intensity measurements). In this case, the FIM predicted variance is much larger than for the other cases (note the change in scales in Fig 7B (bottom right)), but the posterior found by the MH analysis is clearly non-Gaussian. As we will see in the next section, the reason for this failure is likely that the (0,300) min experiment design with this distortion provides insufficient information to identify the model.

#### 3.2.4 PDO-corrected FIM analysis accurately ranks designs for most informative transcription-repression experiment

To demonstrate the practical use of FIM for experiment design, we next asked what design for a Trp-based transcription repression experiment would be best to improve our model of mRNA expression identified in Fig 7. We restricted the set of possible experiments to the previous data set (135 cells at *t*_1_=0 min and 62 cells at *t*_2_=300 min) plus an additional set of 100 cells at a new time *t*_3_ after Trp application, where *t*_3_ could be any time in the allowable set *t*_3_ ∈ [0, 6, 12, 18, …, 1200] min.

We drew 20 random parameter samples from the previous 20,000-sample MH chains that were estimated using data at *t*_1_=0 and *t*_2_=300 min for each data type. We computed the FIM for each parameter set and for every potential choice for *t*_3_. Figure 8A shows the determinant of the expected covariance of MLE parameters (Σ ≈ **FIM**^−1^, defined in log10 parameter space) versus *t*_3_ assuming direct observation of smiFISH mRNA (left) or distorted observations of MCP-GFP spots, FISH Intensity, or GFP intensity (right). As expected, distortion always increases expected uncertainty (compare Figs 8A(left) to the other columns). We also find that the optimal time for the experiment can be highly dependent on the particular assay. In particular, larger distortions require earlier sampling times to prevent mRNA expression from falling below the noise introduced by the distortion.

**Figure 8:**
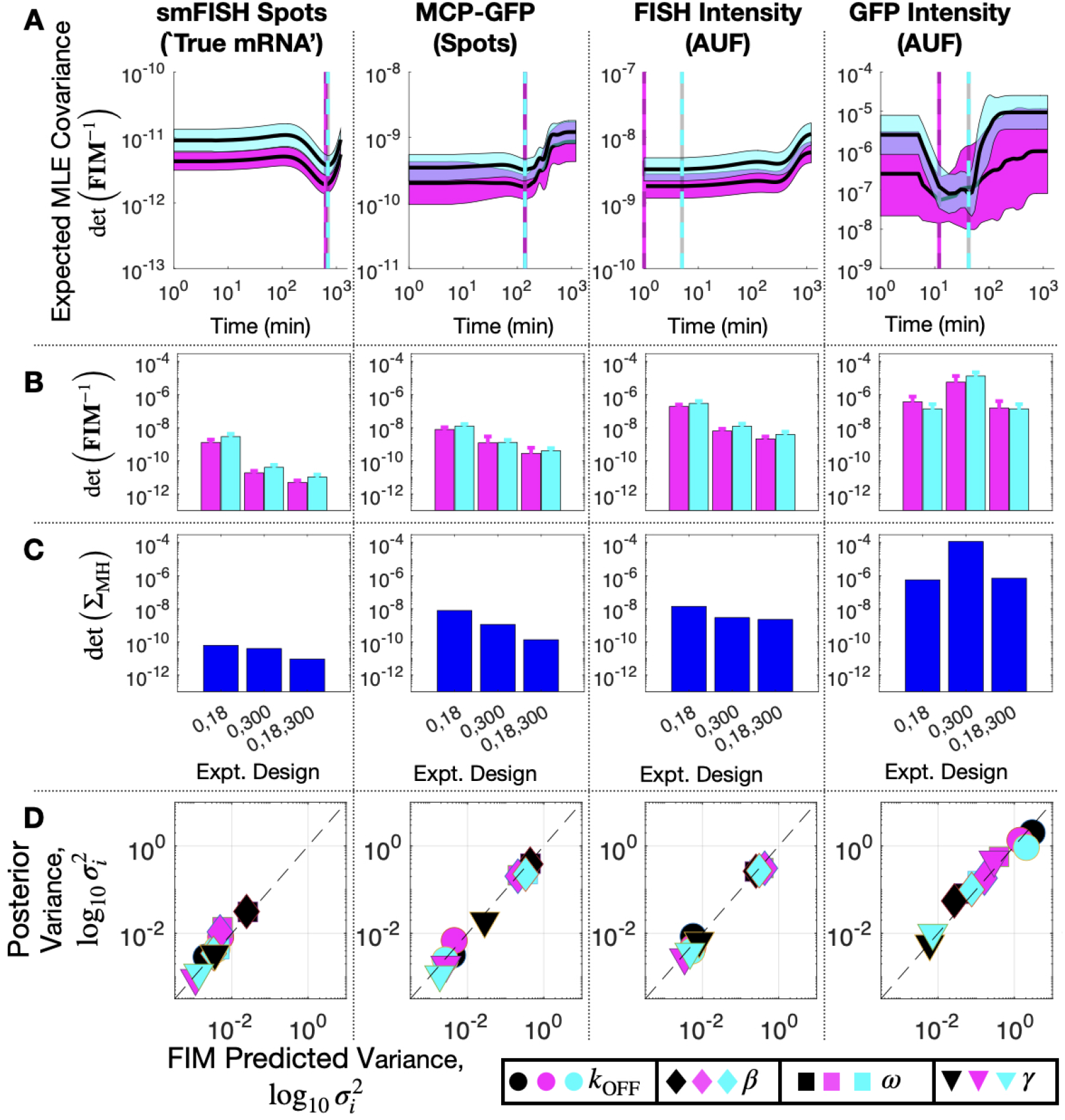
Design of Subsequent Experiment for MS2X128 cassettetagged HIV-1 reporter gene. **(A)** Expected volume of uncertainty (det(**FIM**^−1^)) versus time of third measurement assuming 100 cells and measurement of: (left to right) smiFISH mRNA, MCP-GFP spots, FISH intensity, or GFP intensity. Solid lines and shading denote mean ±SD for 20 parameter sets selected from MH chains after fitting initial data (magenta, *t* =(0,300) min) or final data (cyan, *t* =(0,18,300) min). Cyan and magenta vertical lines denote the optimal design for the third experiment time assuming the corresponding parameter values. **(B)** Expected volume of MLE uncertainty (det(**FIM**^∡1^)) for different sets of experiment times and measurement modalities and averaged over 20 parameters sets sampled from the MH chains for initial fit (magenta) or final parameter estimates (cyan). **(C)** Volume of MLE uncertainty (det(Σ_MH_) estimated from MH analysis in the same experiment designs as **B. (D)** Posterior variance versus FIM prediction of variance for each parameter (symbol key at bottom right), for each measurement modality (different columns) and for analyses based on different sets of data: *t* =(0,18) min (black), *t* =(0,300) min (magenta), or *t* =(0,18,300) min (cyan). All MH analyses contain 20,000 samples. Measurements include 135 cells at *t* = 0, 96 at *t* = 18, and 62 at *t* = 300. Parameter uncertainties defined in log base 10 for all panels.

Although a complete validation of these designs would require a host of experiments that are beyond the scope of the current investigation, we analyze a new data set of 96 cells taken at *t*_3_ = 18 min to compare experiment designs with time combinations of (0,18) min, (0,300) min, and (0,18,300) min. Figure 8B shows FIM predictions of uncertainty using parameters from the original fit to the (0,300) min data (magenta) compared to the uncertainty estimate using parameters fit to the final data set with all three time points (cyan). The original and final FIM are in agreement for all combinations of experiment designs (different groups of magenta and cyan bars) and for direct observations of smiFISH mRNA (left) or for any of the distorted data sets. Moreover, upon using the MH to estimate the posterior parameter distributions, Fig 8C shows that both the original and the final FIM correctly predict the trend of parameter uncertainties for each of the different experiment combinations. For example, the FIM correctly predicts the result that a set of (135,62) cells at (0,300) min is more informative than the larger set of (135,96) cells at (0,18) min if one uses the undistorted data (far left), but the opposite is true if one considers the distortion due to measurements of GFP intensity (far right). Finally, Fig 8D shows the variance in each parameter’s estimation for every measurement type as predicted by the FIM analysis (horizontal axis) and verified using 20,000 MH samples of the posterior (vertical axis). The results show a clear correlation between the predicted and measured uncertainty under either the initial (t=0,300, magenta) or the final (t=0,18,300, cyan) data sets. The analysis also correctly predicts which parameters are well-identified under which measurements. For example, the FIM analysis correctly predicts that the degradation rate *γ* (triangles) is well identified using just t=0,300 min data for the undistorted data, the MCP-GFP spot data, or the FISH intensity data, but requires the t=18 min data to be identified using the GFP intensity data. In other words, the FIM analysis now explains the failure to identify well-constrained parameters that we observed at the end of the previous section, and correctly suggests that this failure can be substantially ameliorated with an additional measurement of at 62 cells at 18 min.

## 4 Discussion

Parameter estimation is a major step in constructing quantitative models for all physical or biological processes, and many such models for gene regulation and cell signaling are now being inferred from quantitative single-cell imaging experiments. Such measurements are subject to errors, where different labels or different image processing can yield different measurement values (e.g., see Figs 5 and 6). We have demonstrated that these experimental distortions can be mathematically described using a general class of probability transition kernels that we dub *Probability Distortion Operators* (PDOs, Figs 2,5B,E), and that these PDOs lead to changes in the estimated parameters that are reflected not only in the magnitudes but also in the directions of their uncertainties (Fig 4, 7 and 8). We have introduced a new computational framework to systematically account for these noise effects and provide a first step toward integrating PDOs into the interpretation of single-cell experiments (Figs 5-8). Our results indicate that an appropriate statistical analysis coupled with a careful tuning of experimental design variables can meaningfully compensate for measurement noise in the data. For example, our results indicate that, when used iteratively with small sets of experimental data (e.g., less than a couple hundred cells at only two points in time), FIM analysis can correctly predict which subsequent experiments are most likely to be informative, and which are unlikely to provide additional insight into model parameters (Fig 8). Insight provided by such an integration of models and experiments could allow for better allocation of experimental resources first by helping to eliminate estimation biases that are due to experimental noise and second by helping to identify specific experimental conditions that are less prone to be impacted by those measurement artifacts.

Although the FIM is a classical tool for optimal experiment design that has been used extensively in myriad areas of science and engineering ([63, 18, 13, 69, 12, 89]), it has not seen widespread adoption in the investigation of biological processes, in part because biological processes are heavily subject to heterogeneities that are not accounted for in traditional FIM analyses. However, there has been some progress to extend these tools to the context of gene expression modeling; for example, ([42]) proposes a method to approximate the FIM for single-cell experiment data based on the Linear Noise Approximation (LNA). Alternatively, the FIM has been approximated by using moment closure techniques ([69, 70]). These approaches work well in the case of high or moderate molecular copy numbers, but they break down when applied to systems with low molecular copy numbers ([23]), and it is not clear how or if such approximations can be modified to consider measurement noise and data processing noise that are non-additive, asymmetric, or non-Gaussian as is the case for many biological distortions. To circumvent these issues, our alternative framework directly analyses the probability distributions of the noisy measurements. Explicitly modeling the conditional probability of the observation given the true cell state allows us to express the observation distribution as a linear transformation of the true process distribution that is computable using the finite state projection. This leads to a systematic way to develop composite experimental designs that combine measurements at different fidelity and throughput levels to maximize information given a budgetary constraint.

The current investigation provides a few examples and preliminary experimental data to illustrate the broad potential of the FIM and PDO formulations to improve the interpretation and design of single-cell experiments. However, these are limited at present, and there is much that remains to be done to elucidate the full capabilities for these new techniques. For example, not only is additional experimental testing needed to validate the use of FIM-based methods for a broader range of experiment designs and imaging conditions, but the current analyses also need further development (i) to allow for more complex definitions of PDOs, (ii) to expand the use of FIM and PDO analyses to situations where prior knowledge and models are unavailable or limited, and (iii) to extend the FIM analyses for use in important tasks of model reduction or model selection. A few of these limitations and future directions are discussed as follows.

The current investigation uses high- and low-fidelity calibration experiments to parameterize three different PDOs for different measurement modalities, selects the PDO that minimizes the Bayesian Information Criteria in each case, and confirms that this selection also led to the best prediction of held out data. However, our search over possible PDOs was far from exhaustive, and it is almost certain that more accurate PDOs could be found. Adjustments are easily made so that PDOs can capture many different aspects of experimental error or so that they can be applied to many different types of models. For example, through experimental analysis of mRNA counts using different modalities (e.g., smiFISH and MCP-GFP labeling), we demonstrated how one could construct the PDO based on empirical measurements (Fig 8). Similarly, one could examine different microscopes, different cameras, different laser intensities, different image processing pipelines, or any of a number of permutations to compare and quantify differences between high-fidelity and noisy measurements of signaling or gene expression phenomena. In the current work, we have relied on specific parameterized statistical distributions (e.g., binomial or Poisson as examined in Figs 3,4,5 or data binning or categorization in Section S3.2) or mechanistic distortion models (e.g., integrated fluorescence intensity in Section S3.1, stochastic binding kinetics in Section S3.3, or segmentation errors in Section S3.4) to formulate the PDO. However, one could also envision parameter-free statistical methods based on recent probabilistic machine learning methods (e.g., normalizing flows ([41]) for modeling the conditional distribution *p*(*y*|*x*) of a noisy label *y* given a vector *x* of features. In either case, one would need only to calibrate the PDO once for each combination of labeling, microscopy, and imaging techniques and then one could to apply that PDO to many different biological processes, models, parameter sets or experiment designs. For simplicity, we have assumed that calibration data is available and that the PDO is constant in time. However, the formulations of the PDO and FIM (Eqs. 3–7) are sufficiently general such that these requirements can be relaxed. With these relaxations, the FIM could be used either to guide experiment designs to aid the simultaneous identification of both a parameterized PDO and the gene regulation model itself, or provide clear guidance that such distortions lead to degeneracy in the FIM and indicate which non-identifiable parameter combinations (i.e., the null space of the distorted FIM) are most in need of model reduction.

An important limitation of any model-based experiment design approach is that to make predictions, one must have some prior knowledge about the system under investigation. In the experimental example above, we used insight gathered in a previous biological context (i.e., live-cell analyses of nascent nascent transcription sites from [21]) to guess some model parameters (i.e., the transcription burst sizes and frequencies), and we used general knowledge of the cell line to guess other parameters (e.g., the half-life for mammalian mRNA). For many single-cell optical microscopy investigations, such information is available in advance, due to the fact that one must choose which genes or pathways to investigate before designing smFISH probes or modifying cells and promoters to express the MS2-MCP reporters, and this choice is typically based on experience or previous investigations in the literature (e.g., on analyses of related genes or pathways, in different environmental conditions, or with other exploratory experimental techniques). However, for earlier stage exploratory investigations, where such prior knowledge may not be available, one may need to collect some preliminary data before building models (e.g., collect data for a small handful of time points). In this case, computing the FIM after fitting to the first round of experiments can help to elucidate which parameter sets of the models are well-identified (i.e., vectors in parameter space corresponding to large eigenvalues of the FIM after the *initial* experiment), which can be improved with different experiment designs (i.e., vectors in parameter space that correspond to larger eigenvalues of the FIM for *different* experiments), and which cannot be identified for any experiment (i.e., vectors in parameter space that lie in the null space of the FIM no matter what experiment is considered).

Although we have only considered one model in the main text for the current investigation, the insight provided by the FIM could also be utilized to analyze multiple candidate model structures, an important task that has previously been explored under the assumption that smFISH yields exact measurements of mRNA content [59, 37, 40]. As discussed above, the FIM is useful to identify and prune highly-uncertain parameter combinations or remove unidentifiable model mechanisms. For example, using the distorted FISH intensity measurements at *t*=(0,300), the FIM analysis clearly shows that the burst frequency (log_10_ *ω*) and burst size (log_10_ *β*) are jointly uncertain along the negative diagonal (Fig 7B, rightmost column), meaning that the average total production rate and standard deviation (*ωβ ≈* 1.04 ± 0.16*s*^−1^) could be well constrained, while the individual parameters *β* = 5.0 ± 5.8 and *ω* = 1.1 ± 2.8 could not be independently identified. A similar observation holds using the MCP-GFP spot count data. In both cases, without the constraint of the prior, this uncertainty would extend to the limit where *β* is much less than one, at which point the proposed 3-state model with parameters [*k*_ON_, *k*_OFF_, *ω, β, γ*] reduces to an equivalent 2-state model with parameters [*k*_ON_, *k*_OFF_, *k*_r_ *≈ ωβ, γ*]. Indeed, Fig S11 shows that this simpler model (Fig S11A) provides nearly (but not quite) as good fits to the all data types when estimated from the *t*=(0,300) min data (Fig S11B, left columns, compare fit likelihood and BIC values to Fig 7B), but these fits have much less parameter uncertainty (Fig S11B, right columns). Moreover, for the smiFISH mRNA counts, the MCP-GFP spot counts and the MCP-GFP intensity data, the simpler model led to *better* predictions for the held out data at *t* =18 min (Fig S11B, middle columns compare prediction likelihood values to Fig 7B). Moving forward, if one’s goal were to differentiate further between these or other competing hypotheses for model mechanisms, one could use FIM-based experiment design to suggest conditions that promise strong uncertainty reduction for several competing models at once. For example, Fig S12 shows the variance reduction predicted for various possible experiment designs for the reduced model. Comparing FIM analyses of potential experiment designs to the results to the original model in Fig 8, we find that for all distortions, experiments that reduce uncertainty for one model should also be effective for the other. However, the purpose of the current study is only to introduce the FIM+PDO formulation, and a complete analysis of the use of FIM insight for model selection is left for future investigations.

Whether one starts with initial parameter and model structures guesses from previous experimentation or based on a preliminary round of experiments, subsequent model identification is most effective when pursued as an iterative endeavor, requiring evaluation of uncertainty and model-driven experiment design at each stage. For example, in our analysis of the HIV-1 reporter gene, initial data at 0 and 300 min after Tpt revealed that the practical identifiability of parameters depended heavily on which measurement was used. Assuming ideal measurements (i.e., using smiFISH), Fig 8B shows that the model with (*n*=4) free parameters could be identified to an log10-uncertainty volume of det(**FIM**^−1^) = 1.88 × 10^−11^ if we include the prior or det(**FIM**^−1^) = 1.91 × 10^−11^ if we do not include the prior. However, using total FISH intensity, the model was much less certain at det(**FIM**^−1^) = 6.39 × 10^−9^ with the prior or det(**FIM**^−1^) = 1.43 × 10^−6^ without the prior. Since the determinant of the *n*-parameter FIM scales with the number of cells according to: det(*α***FIM**) = *α*^*n*^ det(**FIM**), it would take a factor of *α* = (1.43 × 10^−6^/1.91 × 10^−11^)^1/4^ = 16.6 times as many cells using FISH-intensity to achieve the same accuracy as with the smiFISH mRNA data.^1^ Moreover, we also see that the optimal subsequent experiment design also depends heavily on which measurement modality is used. For smiFISH experiments that are assumed to be free from observation noise, the optimal next time point is very late (t=696 min), whereas for the distorted observations, measurements should be taken much earlier (e.g., at *t*=138 min for MCP-GFP spots, at *t*=0 min for FISH intensity, and at *t*=42 min for GFP intensity). Furthermore, in the worse case, choosing the next experiment based on an incorrect assumption for the PDO could lead to waste of experimental efforts – e.g., using the long time as suggested by the smiFISH analysis would be almost entirely worthless if used with one of the other measurements assays. The current study, which is meant only to introduce the FIM and its use for experiment design, has limited availability of experimental data (i.e., one replica with three time points) and only for an artificial HIV-1 reporter construct. A full examination for the use of FIM in iterative single-cell experiment design for endogenous gene regulatory pathways is ongoing and will be described in future publications.

For the model under consideration to fit the HIV-1 promoter with the 128XMS2 stem-loop cassette, computing the CME solution took an average of 1.3s in Matlab on a 2019 MacBook Pro (2.6 GHz 6-Core Intel Core i7), computing the FIM took an average of 40s (39.94s to solve the sensitivity to all parameters and 0.06s to compute the FIM), and running the MH for 20,000 samples took 26,000s, thus the FIM estimates uncertainty roughly 650 times faster than the MH. For use in experiment design, the cost savings provided by the FIM can be much higher. Because one can reuse pre-computed sensitivities, the computational cost is only 0.06s for each new experiment design (e.g., to explore different numbers of cells, different PDOs, or differe data (e.g., to generate MLE scatter plots as shown Figs 3C, 3D or S5), if one liberally assumed that an experiment could be evaluated using only *N*_*s*_ =100 samples and that adequate fits could be achieved using only *N*_*f*_ =100 parameter guesses (usually far more function evaluations are needed), then evaluating each new experiment design would require one CME solution (1.3s) to generate data and 1.3 *· N*_*s*_ *· N*_*f*_ = 13,000s to fit those data (217,000 times longer than the FIM approach). Given a priori uncertainty in the model, in practice one would need to redo both the FIM analysis (40s + 0.6s per experiment) or the simulation-based analysis (13,000s per experiment) for many different parameter sets or model structures (e.g., we show results from 20 parameter combinations in Figs 8A and S12A), and the savings provided by the FIM becomes even more important.

Finally, although the presented approach is versatile and can in principle be applied to any stochastic gene regulatory network, its practical use depends on the ability to compute a reasonable approximation to the solution of the chemical master equation (CME) as well as its partial derivatives with respect to model parameters. Fortunately, there are now many relevant stochastic gene expression models for which exact or approximate analytical expressions for the CME solution are available ([61, 77, 85, 33, 10, 90]). Furthermore, the FSP and similar approaches have been used successfully to solve the CME for many non-linear and time-inhomogeneous regulatory models for which closed-form solutions do not exist ([80, 15, 51, 59, 79, 75]). For example, SI Section S4 analyzes a model of the nonlinear genetic toggle switch, and the FIM and PDO analysis is used to ask which species should be measured and for how many cells in order to best identify model parameters. As another example, SI Section S5 analyzes a spatial stochastic model with a four-state gene expression model under time varying MAPK activation signal and nucleus to cytoplasmic transport[57]. Admittedly, given the complexity of gene regulatory networks in single cells, there will always be stochastic gene regulatory models whose direct CME solutions are beyond reach. Nevertheless, continued advancements in computational algorithms ([38, 9, 29, 49, 11, 30, 60]) are enlarging the set of tractable CME models, which in turns can help accelerate the cycling between data acquisition, model identification, and optimal experiment design in single-cell studies.

## Author Contributions

Conceptualization: BM. Theory and computational modeling: HDV and BM. Performed experiments/collected data: LSFQ. Image processing: LU. Writing: HDV, LSFQ, LUA, and BM. Resources, supervision, and funding acquisition: BM.

## Acknowledgments

We thank members of Munsky group for valuable comments on the manuscript, especially Eric Ron for suggesting the investigation of cell segmentation noise.

## Data Availability Statement

Python code for the simulation and FIM analysis of the two-state bursting gene expression model is available at:

https://github.com/voduchuy/NoisyMeasurementFIM

Python code for processing single-cell images is available at:

https://github.com/MunskyGroup/FISH_Processing

Matlab code for analyzing Tryptolide data using the three-state model is available at:

https://github.com/MunskyGroup/SSIT/, where all data and codes for fitting and visualization of results are available in the folder “CommandLine/Vo et al 2023”.

## Footnotes

1 We note that total intensity is much easier to compute than spot detection and could in principle be measured at lower microscope resolution or even with flow cytometry. Depending upon available equipment, collecting 16.6 times as many cells could potentially be achieved at a lower overall experimental cost!

