## Supporting Information for "Analysis and design of single-cell experiments to harvest fluctuation information while rejecting measurement noise"

April 25, 2023

#### Contents

**S1 Overview of the Chemical Master Equation and the Finite State Projection S1**

**S2 Corrected and uncorrected likelihoods for fitting noisy datasets S3**

**S3 Additional information and examples for probabilistic distortions applied to the random telegraph model S3**

S3.1 Numerical computation of measurement-distorted single-cell mRNA count and intensity distributions . . . S4
S3.2 Effect of binning observation data on information and optimal experiments . . . S5
S3.3 Defining the PDO using a second CME to account for stochastic dynamics in the experimental measurement assay . . . S6
S3.4 Effects of image segmentation distortions on information and optimal experiment design . . . S8

**S4 Two-species toggle-switch case study: Optimal combinations of different measurement modalities can provide tremendous reductions to parameter uncertainties S11**

**S5 Exploring the effects of partial or lumped observations for the identification of spatially-compartmentalized gene expression models S13**

#### S1 Overview of the Chemical Master Equation and the Finite State Projection

Molecular fluctuations in single cells can be modeled using stochastic reaction networks (SRNs) [1]. This is a stochastic, discrete extension of the classical reaction rate description. Consider a system in the cell with

*Department of Chemical and Biological Engineering, Colorado State University
†Department of Chemical and Biological Engineering, Colorado State University
‡Department of Chemical and Biological Engineering, Colorado State University
§Department of Chemical and Biological Engineering, School of Biomedical Engineering, Colorado State University

$n_S$ chemical species that can interact through $n_R$ reaction channels. The molecular counts of these species can be collected into a state vector $\boldsymbol{x} = (x_1, \ldots, x_{n_S})$. The state space of the model consists of all $n_S$-tuples of non-negative integers, representing all possible combinations of molecular counts that a single cell may achieve. The stoichiometry vectors $\boldsymbol{\nu}_j, j = 1, \ldots, n_R$ are defined as the vector of changes to the molecular counts after a reaction event. This means that a cell with state vector $\boldsymbol{x}$ will transit to state $\boldsymbol{x} + \boldsymbol{\nu}_j$ after a firing of reaction $j$.

The SRN framework assumes that the molecular count vector $\boldsymbol{X}(t)$ is a Markov process, whose stochastic dynamics is determined by the propensity functions $\alpha_j(t, \boldsymbol{x}; \boldsymbol{\theta}), j = 1, \ldots, n_R$ that may depend on the time $t$, state $\boldsymbol{x}$, and a vector $\boldsymbol{\theta}$ of model parameters. Intuitively speaking, if the cell is at state $\boldsymbol{x}$ at time $t$, then $\alpha_j(t, \boldsymbol{x}; \boldsymbol{\theta})\mathrm{d}t$ is the probability for the $j$th reaction to occur during the next infinitesimal interval $[t, t + \mathrm{d}t]$.

Let there be an enumeration of the state space so that we can arrange states into a sequence $\boldsymbol{x}_1, \boldsymbol{x}_2, \ldots$. For example, we could apply the Cantor pairing function [5][10] that defines a mapping $\boldsymbol{x} = (x_1, \ldots, x_{n_S}) \mapsto \Phi_{n_S}(\boldsymbol{x})$ via the recursion

$$\begin{aligned}
\Phi_2(x_1, x_2) &= \frac{1}{2}(x_1 + x_2)(x_1 + x_2 - 1) + x_2, \\
\Phi_n(x_1, \ldots, x_{n_S}) &= \Phi_2\left(\Phi_{n-1}(x_1, \ldots, x_{n_S - 1}), x_{n_S}\right).
\end{aligned}$$

The probability distribution of $\boldsymbol{X}(t)$ can be thought of as an infinite-length vector $\boldsymbol{p}(t)$ indexed by such enumeration, with the $i^{\text{th}}$ entry $p_i(t)$ equal to the probability that $\boldsymbol{X}(t) = \boldsymbol{x}$. If we assume that the initial distribution $\boldsymbol{p}(0) = \boldsymbol{p}_0$ is known, then we can, in theory, obtain the time-varying probability distribution $\boldsymbol{p}(t)$ by solving the Chemical Master Equation (CME):

$$\begin{aligned}
\frac{d}{dt}\boldsymbol{p}(t, \boldsymbol{\theta}) &= \boldsymbol{A}(t, \boldsymbol{\theta})\boldsymbol{p}(t, \boldsymbol{\theta}), \\
\boldsymbol{p}(0, \boldsymbol{\theta}) &= \boldsymbol{p}_0(\boldsymbol{\theta}).
\end{aligned}$$

The matrix $\boldsymbol{A}(t, \boldsymbol{\theta}) = [a_{ij}(t, \boldsymbol{\theta})]$ is called the transition rate matrix or infinitesimal generator matrix. Entry-wise, it is defined as:

$$a_{ij}(t, \boldsymbol{\theta}) = \begin{cases} \alpha_k(t, \boldsymbol{x}_j, \boldsymbol{\theta}) & \text{if } \boldsymbol{x}_i = \boldsymbol{x}_j + \boldsymbol{\nu}_k \text{ for some } k, \\ -\sum_{k=1}^{n_R} \alpha_k(t, \boldsymbol{x}_j, \boldsymbol{\theta}) & \text{if } i = j, \\ 0 & \text{otherwise.} \end{cases}.$$

The Finite State Projection (FSP)[11] is based on specifying a finite truncation $\boldsymbol{p}^{\mathrm{FSP}}(t) \in \mathbb{R}^{|J|}$ of the probability vector $\boldsymbol{p}(t)$ by keeping only a finite set of states $J = \{\boldsymbol{x}_1, \ldots, \boldsymbol{x}_{|J|}\}$. A similar truncation is applied to the transition rate matrix $\boldsymbol{A}(t, \boldsymbol{\theta})$ and the resulting finite system of linear ODEs,

$$\frac{d}{dt}\boldsymbol{p}^{\mathrm{FSP}}(t, \boldsymbol{\theta}) = \boldsymbol{A}^{\mathrm{FSP}}(t, \boldsymbol{\theta})\boldsymbol{p}^{\mathrm{FSP}}(t, \boldsymbol{\theta}), \tag{1}$$

can be solved numerically using conventional numerical ODE integration methods. The FSP has a simple and computable a posteriori error estimate [11]

$$\|\boldsymbol{p}(t, \boldsymbol{\theta}) - \tilde{\boldsymbol{p}}^{\mathrm{FSP}}(t, \boldsymbol{\theta})\|_1 = 1 - \sum_{j=1}^{|J|} p_j^{FSP}(t, \boldsymbol{\theta})$$

where $\tilde{\boldsymbol{p}}^{\text{FSP}}$ is the "lifted" version of $\boldsymbol{p}^{\text{FSP}}$ by padding zeros as appropriate for mathematical consistency. In practice, this means that the error is kept below $\varepsilon$ if the total sum of entries of $\boldsymbol{p}^{\text{FSP}}(t;\boldsymbol{\theta})$ is above $1-\varepsilon$.

The FIM analysis in the main text also requires approximations to the sensitivity vectors $\boldsymbol{s}_j(t;\boldsymbol{\theta}) = \frac{\partial}{\partial \theta_j}\boldsymbol{p}(t,\boldsymbol{\theta})$. Following [3], we provide a finite truncation approximation $\boldsymbol{s}_j^{\text{FSP}}$ to each of these sensitivity vectors by solving the forward sensitivity ODEs

$$\frac{d}{dt}\boldsymbol{s}_j^{\text{FSP}}(t,\boldsymbol{\theta}) = \frac{\partial}{\partial \theta_j}\boldsymbol{A}^{\text{FSP}}(t,\boldsymbol{\theta})\boldsymbol{p}^{\text{FSP}}(t,\boldsymbol{\theta}) + \boldsymbol{A}^{\text{FSP}}(t,\boldsymbol{\theta})\boldsymbol{s}_j^{\text{FSP}}(t,\boldsymbol{\theta}). \tag{2}$$

### S2 Corrected and uncorrected likelihoods for fitting noisy datasets

Consider a dataset $\mathcal{D} = \{(t_i, \boldsymbol{y}_i), i = 1, \ldots, n_D\}$ that consists of $n_D$ independent single-cell observations, where the $i^{\text{th}}$ observation is measured at time $t_i$ and the observed expression is $\boldsymbol{y}_i$.

If we do not acknowledge the existence of measurement noise, then it follows that $\boldsymbol{y}_i$ is the true molecular counts in the cells. The likelihood of observing $\mathcal{D}$ given a CME model with parameters $\boldsymbol{\theta}$ is then

$$L^{\text{uncorrected}}(\mathcal{D}|\boldsymbol{\theta}) = \prod_{i=1}^{n_D} p_X(t_i, \boldsymbol{y}_i, \boldsymbol{\theta}), \tag{3}$$

where $p_X(t,\boldsymbol{y},\boldsymbol{\theta})$ is the probability that the *true* cell state $\boldsymbol{X}(t)$ equals to $\boldsymbol{y}$ at time $t$. These probabilities can be computed with the FSP.

On the other hand, if we know that each $\boldsymbol{y}_i$ is a distortion of the true cell state at time $t_i$, its probability given a model is actually

$$p_Y(t,\boldsymbol{y}_i,\boldsymbol{\theta}) = \sum_{\boldsymbol{x}} p(\boldsymbol{y}_i|\boldsymbol{x},\boldsymbol{\theta})p_X(t,\boldsymbol{x},\boldsymbol{\theta}), \tag{4}$$

where $p(\boldsymbol{y}_i|\boldsymbol{x},\boldsymbol{\theta})$ is the probability of observing $\boldsymbol{y}_i$ given the true cell state $\boldsymbol{x}$. This results in the correct likelihood formulation

$$L(\boldsymbol{\theta}|\mathcal{D}) = \prod_{i=1}^{n_D} p_Y(t_i,\boldsymbol{y}_i,\boldsymbol{\theta}). \tag{5}$$

For Fig. 3 in the main text, we generated distorted datasets and use numerical optimization to find the MLE fits to these datasets. The MLE-PDO fits use the correct likelihood formulation (5) as the objective function whereas the MLE fits with uncorrected likelihood use the uncorrected formulation (4).

To locate the MLE fit for each simulated dataset, we use the compass search algorithm [9] implemented in the PyGMO optimization package [2]. Since we want to simulate the effect of data sampling noise on MLE, each numerical search starts from the true data-generating parameters. Due to sampling noise, the actual MLE fits to simulated datasets will differ from the actual data-generating parameters, and it is the spread of the MLE fits that the inverse of the FIM can capture in the asymptotic limit where $n_D \to \infty$ [13].

### S3 Additional information and examples for probabilistic distortions applied to the random telegraph model

In this section, we provide additional examples for using the extended Fisher Information Matrix analysis with different probabilistic distortion operators, assuming the gene expression model of interest is the telegraph model in the main text. As in the main text, we assume an experiment design in which five batches of independent single-cell measurements, each of which consists of $1,000$ cells, are placed at five equally spaced times. The design variable is the sampling period $\Delta t$, which is the time between two successive batch measurements, and we are interested in how either the value of $\Delta t$ or the parameters of the measurement distortion (or both) influences experimental design criteria such as the determinant of the Fisher Information Matrix.

#### S3.1 Numerical computation of measurement-distorted single-cell mRNA count and intensity distributions

For the examples in the main text whose observation variable is in a discrete domain, we apply a truncated approximation similar to the FSP. As in previous sections, let $\boldsymbol{C}(t;\boldsymbol{\theta})$ denote the PDO and $\boldsymbol{p}_X(t,\boldsymbol{\theta})$, $\boldsymbol{p}_Y(t,\boldsymbol{\theta})$ denote the time- and parameter-dependent distributions of the true single-cell state $\boldsymbol{X}(t)$ and observed state $\boldsymbol{Y}(t)$. Let $J_X$ be a subset of the state space for true single-cell states and $J_Y$ be a subset of the space of all observable states. FSP-like approximations for $\boldsymbol{p}_Y(t)$ can be obtained by

$$\hat{\boldsymbol{p}}_Y(t,\boldsymbol{\theta}) = \hat{\boldsymbol{C}}(t,\boldsymbol{\theta})\boldsymbol{p}_X^{\mathrm{FSP}}(t,\boldsymbol{\theta}), \tag{6}$$

where $\boldsymbol{C}^{\mathrm{FSP}}$ is the submatrix of $\boldsymbol{C}(t,\boldsymbol{\theta})$ consisting of rows in $J_Y$ and columns in $J_X$. See Fig 2 in the main text for visualizations of these submatrices for the distortion models of bursting gene mRNA counts in the main text. In all examples in the main text, we let $J_Y = \{0,\ldots,400\}$, and $J_X$ is the state subset selected by the adaptive FSP implementation [16] such that the $\ell$-1 error is below $10^{-8}$. Similarly, we obtain a finite state approximation $\hat{\boldsymbol{s}}_j^Y(t;\boldsymbol{\theta})$ for each of the sensitivity vectors $\boldsymbol{s}_j^Y(t;\boldsymbol{\theta})$ as

$$\hat{\boldsymbol{s}}_j^Y(t,\boldsymbol{\theta}) = \hat{\boldsymbol{C}}(t,\boldsymbol{\theta})\hat{\boldsymbol{s}}_j^X(t,\boldsymbol{\theta}) + \frac{\partial}{\partial\theta_j}\hat{\boldsymbol{C}}(t,\boldsymbol{\theta})\boldsymbol{p}_X^{\mathrm{FSP}}(t;\boldsymbol{\theta}). \tag{7}$$

For the main text example of integrated intensity measurements, the observation space is continuous. The $(i,j)$ element of the FIM is given by

$$F_{ij}(t,\boldsymbol{\theta}) = \int_{\mathcal{Y}} d\boldsymbol{y} p_Y(t,\boldsymbol{y},\boldsymbol{\theta})\frac{s_i^Y(t,\boldsymbol{y},\boldsymbol{\theta})s_j^Y(t,\boldsymbol{y},\boldsymbol{\theta})}{p_Y^2(t,\boldsymbol{y},\boldsymbol{\theta})}. \tag{8}$$

While we cannot form a continuous vector in the same way as in discrete observation space, we can still approximate point-wise probability densities $p^Y(t,\boldsymbol{y},\boldsymbol{\theta})$ and their partial derivatives by multiplying each row of $\boldsymbol{C}(t,\boldsymbol{\theta})$ with the FSP solution $\boldsymbol{p}_X^{\mathrm{FSP}}(t,\boldsymbol{\theta})$. That is, we can evaluate

$$\hat{p}_Y(t,\boldsymbol{y},\boldsymbol{\theta}) = \sum_{\boldsymbol{x}\in J_X}\boldsymbol{C}(t,\boldsymbol{\theta})(\boldsymbol{y},\boldsymbol{x})p_X^{\mathrm{FSP}}(t,\boldsymbol{x},\boldsymbol{\theta}), \tag{9}$$

$$\hat{s}_j^Y(t,\boldsymbol{y},\boldsymbol{\theta}) = \sum_{\boldsymbol{x}\in J_X}\frac{\partial}{\partial\theta_j}\boldsymbol{C}(t,\boldsymbol{\theta})(\boldsymbol{y},\boldsymbol{x})p_X^{\mathrm{FSP}}(t,\boldsymbol{x},\boldsymbol{\theta}) + \boldsymbol{C}(t,\boldsymbol{\theta})(\boldsymbol{y},\boldsymbol{x})\hat{s}_j^X(t,\boldsymbol{x},\boldsymbol{\theta}). \tag{10}$$

Combining these point-wise evaluations with Monte Carlo approximation, we obtain

$$\hat{F}_{ij}(t,\boldsymbol{\theta}) = N^{-1}\sum_{k=1}^{N}\frac{\hat{s}_i^Y(t,\boldsymbol{y}_k,\boldsymbol{\theta})\hat{s}_j^Y(t,\boldsymbol{y}_k,\boldsymbol{\theta})}{\hat{p}_Y^2(t,\boldsymbol{y}_k,\boldsymbol{\theta}),} \tag{11}$$

where $\boldsymbol{y}_1,\ldots,\boldsymbol{y}_N$ are i.i.d. samples from the distribution of $\boldsymbol{Y}(t)$, which in our case is obtained by adding Gaussian noise to a scaled version of $\boldsymbol{X}(t)$ that is readily obtained using stochastic simulation [4]. We use $N = 10^5$ in our calculations.

#### S3.2 Effect of binning observation data on information and optimal experiments

Binning (discretization) is a simple and common approach to processing or compressing discrete (continuous) data, and there are many different choices for binning strategies (e.g., choosing the number and borders of bins). The extended FIM analysis presented in the main text is easily adapted to to study the impact of binning strategies on parameter estimation. Here, we consider uniform binning with different choices for bin widths. For a fixed with $w$, the PDO for uniform binning of width $w$ has the form

$$\boldsymbol{C}_{\text{binning}}(y,x) = \mathbf{1}\{y \cdot w \le x < (y+1)\cdot w\} := \begin{cases} 1 \text{ if } x \in [y\cdot w, (y+1)\cdot w) \\ 0 \text{ otherwise} \end{cases} .$$

Fig. S1 shows the effect of different choices of bin width to experiment design. Increasing bin size generally results in a decrease to the determinant of the FIM (Fig. S1A). However, this decline is not necessarily monotonic as the locations of bin edges also plays an important role for the level of information. In addition, the orientation of the uncertainty ellipses (Fig. S1B& C) are both sensitive to the choice of bin size. Further analysis of the optimal binning strategies, including non-uniform binning, is easily formulated using the extended FIM analysis, but an extensive analysis for the interplay between binning strategies and models is beyond the scope of the current study and is left for future investigations.

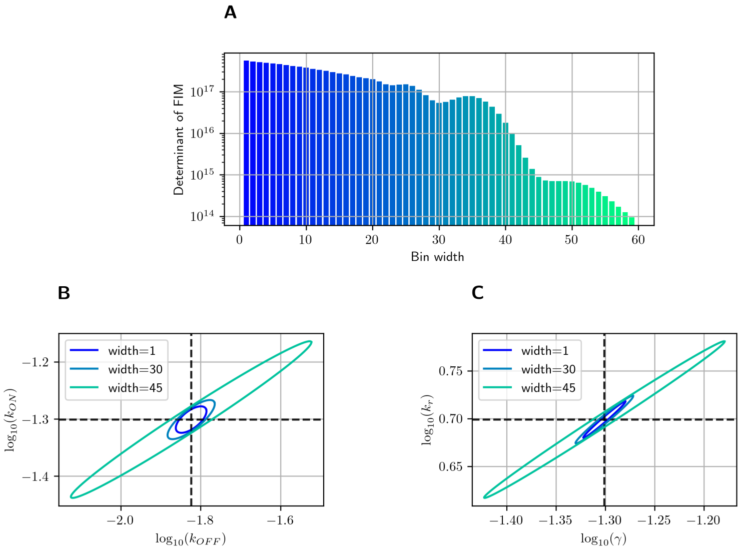


Figure S1: Effect of uniform binning on parameter estimation for the random telegraph model. **(A)**: Determinant of the FIM corresponding to three different choices of bin sizes coupled with different sampling periods. **(B)**: D-optimal information (bar charts, left axis) and sampling periods (solid line, right axis) associated with different choices of bin widths. **(C)**: Three-sigma confidence ellipses on the $\log_{10}(k_{ON}) - \log_{10}(k_{OFF})$ plane associated with three different bin widths. **(D)**: Same as (C) but on the $\log_{10}(k_r) - \log_{10}(\gamma)$. We assume that experimental data comes from collecting five batches of $1,000$ cells each at five uniform sampling times $j\Delta t, j = 1,2,3,4,5$ with $\Delta t := 30$ minute.

#### S3.3 Defining the PDO using a second CME to account for stochastic dynamics in the experimental measurement assay

The single-molecule FISH approach to transcription visualization in fixed cells depends on the binding of fluorescent probes to the mRNA molecules [14]. This binding process is subject to its own molecular fluctuations that control which mRNA are labeled and which are not. Therefore, it can be instructive to model the dynamics of probe binding in an smFISH protocol using another stochastic model informed by the probe's biochemistry. For a simple demonstration, we introduce a toy model here to describe stochastic probe binding to mRNAs (Table S1). This model consists of a second CME separated from the original CME that describe bursting dynamics. The model keeps track of three species: unbound mRNA molecules (hidden to the microscope), mRNA molecules bound to probes (visible to the microscope), and false spots that result from non-specific probe binding or clumping. After exposing the cells to probes for a finite amount of time, the observed number of spots per cell is given by the sum total of mRNA molecules bound by probes as well as false spots. The kinetic parameters of the probe binding/unbinding and the formation of false spots depend on probe concentration. Higher probe concentrations improve the chance of an mRNA molecule to be visualized and detected, but also increase the risk of false positives.

The conditional probability $P(n_{spot}|n_{RNA})$ of observing $n_{spot}$ spots given the true copy number $n_{RNA}$ is computed as following:

1. Initiate the (second) CME solver with initial state $\boldsymbol{z}_0 = (n_{RNA}, 0, 0)$, where the first entry is the copy number of latent RNA, the second is the copy number of visualized RNA, and the third is the number of false spots.

2. Solve the CME using, say, the FSP, up to the maximum time the cells are exposed to the probes $t_{\text{exposure}} := 300$ (AU).

3. From the (second) CME solution, derive the probability distribution of $n_{spot}$ using the formula

$$P(n_{spot}|n_{RNA}) = \sum_{\boldsymbol{z}} (z_2 + z_3) p_{\text{CME}}(t_{\text{exposure}}, \boldsymbol{z}).$$

Collecting these conditional probabilities, we can obtain the associated PDO (up to some truncation). Fig S2 displays four different PDOs associated with four different levels of probe concentrations, which we chose as $0.1, 1.0, 5.0$, and 10.0 arbitrary unit (AU). Observing Fig S2, we can see that with low probe concentration (0.1 AU), there is high risk of missing mRNA molecules. On the other hand, at high probe concentration (10.0 AU), we have significant probability of over-reporting the number of spots due to the formation of false spots. Fig S3 displays how such a trade-off affects the information in experiment design (Fig S3A& B) as well as parameter uncertainties (Fig S3C& D). We can see that probe binding fluctuations reduce information compared to the ideal measurement (Fig S3A& B), and the maximum amount of achievable information is achieved at an intermediate concentration (1.0AU, see Fig S3B). This extension of the FIM analysis to include stochastic dynamics of the measurement assay can similarly be used to account for many different smFISH error sources such as stochastic variations in cell fixation time or variations in probe permeability through cellular membranes. Similarly, the same mathematical analysis can be used to explore effects of RNA dropout or amplification in the analysis of single-cell RNA sequencing data. Such explorations are beyond the scope of this study and are left to future investigations.

| Reaction | Propensity |
|---|---|
| $\mathrm{RNA_{unbound}} \rightarrow \mathrm{RNA}_{bound}$ | $k_{\mathrm{bind}} \cdot \mathcal{C} \cdot [\mathrm{RNA_{unbound}}]$ |
| $\mathrm{RNA_{bound}} \rightarrow \mathrm{RNA}_{unbound}$ | $k_{\mathrm{unbind}} \cdot [\mathrm{RNA_{bound}}]$ |
| $\emptyset \rightarrow$ False spot | $k_{\mathrm{clumping}} \cdot \mathcal{C}$ |
| False spot $\rightarrow \emptyset$ | $k_{\mathrm{dissolution}} \cdot$ [False spot] |

Table S1: Reactions and propensity functions for a second CME model to generate the probabilistic distortion for RNA visualization due to incomplete binding and artificial clumping of smFISH probes. The constant $\mathcal{C}$ is the concentration of probe, in arbitrary units (AU). The other kinetics constants are fixed at: $k_{\mathrm{bind}} := 10^{-1}$, $k_{\mathrm{unbind}} := 10^{-2}$, $k_{\mathrm{clumping}} := 10^{-1}$, $k_{\mathrm{dissolution}} := 5 \times 10^{-2}$ also in arbitrary units.

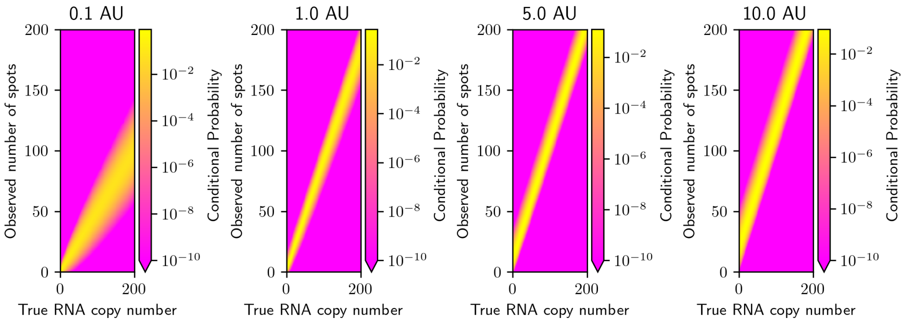


Figure S2: Probabilistic Distortion Operators formulated by a second Chemical Master Equation to describe stochastic fluctuations in probe binding. Propensity functions for the reactions to generate these PDOs are provided in Table S1.

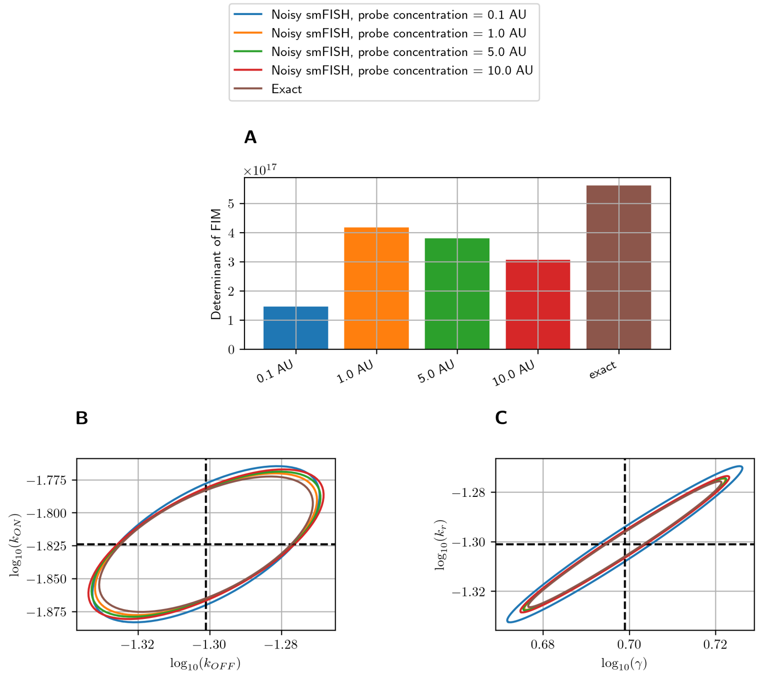


Figure S3: Effect of probe concentration on experiment design for estimating telegraph gene expression parameters. **(A)**: Determinant of the FIM associated with different sampling periods using exact measurements as well as noisy smFISH with four different probe concentrations. **(B)**: D-optimal experiment designs achievable by each probe concentration level (the ideal case of exact measurement is plotted for comparison). **(C)**: Three-sigma confidence ellipses on the $\log_{10}(k_{ON}) - \log_{10}(k_{OFF})$ plane associated with the four probe concentrations and ideal measurements. **(D)**: Same as (C) but on the $\log_{10}(k_r) - \log_{10}(\gamma)$. We assume that experimental data comes from collecting five batches of $1,000$ cells each at five uniform sampling times $j\Delta t, j = 1, 2, 3, 4, 5$ with $\Delta t := 30$ minute.

#### S3.4 Effects of image segmentation distortions on information and optimal experiment design

Cell segmentation is a key step in processing single-cell microscopy images to count single-cell mRNA using smFISH [7]. We consider here a situation that arises when the segmentation algorithm mistakenly groups multiple cells together to classify them as a single object. This could happen, for example, when the cells are crowded and when the signal is low (e.g., if the cells are imaged without a stain for the nucleus or cytoplasmic) or if gates set for flow cytometry analysis permit doublets to be counted as single cells. As a consequence, the molecular count one observes for an object after image processing or cytometry detection may have a chance of actually being the sum of mRNA counts in two or more cells. For simplicity, we assume that one segmented cell may contain only one or two actual cells and that spot counts are exact. However, the analysis can be extended to consider triplets or more cells per object and can also be extended to include any of the distortion effects discussed above or in the main text.

Let $\rho \in [0, 1]$ be the probability that an observed molecular count is actually the sum from two actual cells. The observed molecular copy number $Y$ thus takes the form

$$Y = X + \mathbb{1}\{U \leq \rho\}X_1,$$

where $X$ is the true copy number, distributed according to the solution of the CME, $X_1$ is independent from $X$ but is identically distributed, and $U$ is an independent uniform random number in $[0, 1]$. The distribution of observed counts is related to the distribution of $X$ by

$$\boldsymbol{p}_Y(t, \boldsymbol{\theta}) = (1 - \rho)\boldsymbol{p}_X(t, \boldsymbol{\theta}) + \rho\boldsymbol{p}_X(t, \boldsymbol{\theta}) \star \boldsymbol{p}_X(t, \boldsymbol{\theta}), \tag{12}$$

where $\star$ denotes the convolution operation. The PDO $\boldsymbol{C}(t; \boldsymbol{\theta})$ in this case is the linear map $\boldsymbol{u} \mapsto (1 - \rho)\boldsymbol{u} + \rho\boldsymbol{p}_X(t; \boldsymbol{\theta}) \star \boldsymbol{u}$. We use a finite-dimension approximation to $\boldsymbol{p}_Y(t)$ by substituting the FSP approximation of $\boldsymbol{p}_X$, resulting in the expression

$$\hat{\boldsymbol{p}}_Y(t, \boldsymbol{\theta}) = (1 - \rho)\begin{bmatrix}\boldsymbol{p}_X^{FSP}(t, \boldsymbol{\theta}) \\ \boldsymbol{0}_{n-1}\end{bmatrix} + \rho\text{conv}\left(\boldsymbol{p}_X^{FSP}(t, \boldsymbol{\theta}), \boldsymbol{p}_X^{FSP}(t, \boldsymbol{\theta})\right), \tag{13}$$

where $n$ is the length of the FSP approximation vector $\boldsymbol{p}_X^{FSP}$, $\boldsymbol{0}_{n-1}$ is a vector of zeros of length $n - 1$ for padding, and 'conv' is the vector convolution operation. From this, we also derive finite approximations to the sensitivity vectors by

$$\hat{\boldsymbol{s}}_j^Y(t, \boldsymbol{\theta}) = (1 - \rho)\begin{bmatrix}\boldsymbol{s}_j^{FSP}(t, \boldsymbol{\theta}) \\ \boldsymbol{0}_{n-1}\end{bmatrix} + 2\rho\text{conv}\left(\boldsymbol{p}_X^{FSP}(t, \boldsymbol{\theta}), \boldsymbol{s}_j^{FSP}(t, \boldsymbol{\theta})\right), \tag{14}$$

which, along with $\hat{\boldsymbol{p}}_Y(t, \boldsymbol{\theta})$, allows us to compute an approximation to the FIM of $Y(t)$.

We consider the same experimental setup in the main text, in which five independent batches of $1,000$ cells each are collected at five uniform times $k\Delta t, k = 1, \ldots, 5$, but now with observations distorted by segmentation noise. Fig. S4 displays the FIM-based comparison of different segmentation noise levels in terms of their effects on information in the resulting data (quantified by the determinant of the FIM) and uncertainty estimates.

At first glance, it may seem counter-intuitive that even at the maximal noise level $\rho = 1$, where every "single-cell" count is actually the sum of two independent real cells, there is no loss in information or increase in uncertainty. Indeed, if every measurement contains two cells, then it is possible to extract more information, using the same model. To confirm that this is not an error in our FIM calculation, we perform independent validation using maximum-likelihood fits with simulated data with and without cell aggregation distortions. The result confirms that the computed FIMs indeed provide good estimate of the uncertainties in MLEs (Fig. S5) for the tested case where $\rho = 1$, and the MLE fits to noise-free and distorted datasets are indeed close. The intuitive explanation for these results is that each dataset of $1,000$ noisy observations at $\rho = 1$ is in fact the probabilistically distorted measurement of a larger dataset of $2,000$ noise-free observations (but where pairs of cells are randomly merged together to produce only 1000 observations). When the counts for exactly two cells are always merged together, some aspects of the sampled probability distributions will become easier to estimate (e.g., the standard error of the mean is smaller when a greater number of data points are used to estimate the mean). As a result some combinations of parameters become easier to identify, while others may become more difficult. For example, in Fig. S5(left), identification using singlets only (green ellipse) does a better job to reduce uncertainty along the positive diagonal, while identification using only doublets (red ellipse) does a better job to reduce uncertainty along the negative diagonal. The quantitative effect of cell aggregation depends upon the specific model and experimental conditions, but a full exploration of optimization under cell aggregation distortion is beyond the scope of this study and left for future exploration. However, beyond this intuition to explain how doublet cells may be more informative than singlets, there is more general mathematical observation – if the PDO itself depends on the parameters to be identified, then the distorted measurements can be more informative than the undistorted measurements! Full exploration of this effect is left for future investigations.

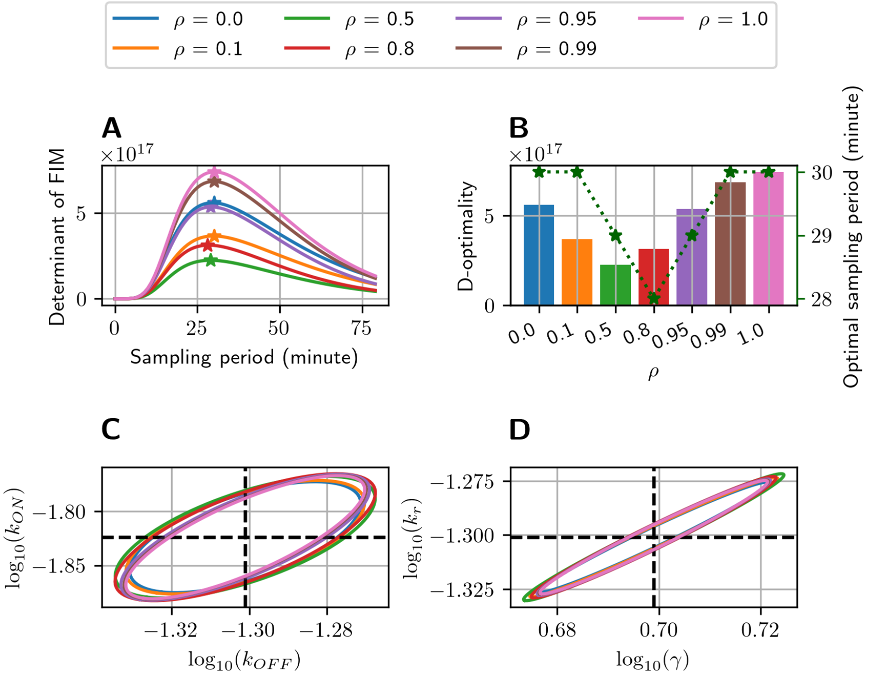


Figure S4: Effect of mis-segmentation on parameter estimation for the random telegraph model (see supplemental text S2.5 for details on the noise model). **(A)**: Determinant of the FIM corresponding to different levels of segmentation noise coupled with different sampling periods. **(B)**: D-optimal information (bar charts, left axis) and sampling periods (solid line, right axis) associated with different segmentation noise levels. **(C)**: Three-sigma confidence ellipses on the $\log_{10}(k_{ON}) - \log_{10}(k_{OFF})$ plane. **(D)**: Same as (C) but on the $\log_{10}(k_r) - \log_{10}(\gamma)$. We assume that experimental data comes from collecting five batches of $1,000$ cells each at five uniform sampling times $j\Delta t, j = 1, 2, 3, 4, 5$ with the sampling period $\Delta t$ varying in panel (A) and fixed at the optimal value associated with the segmentation noise level $\rho$ in panels (B), (C) and (D).

| Reaction | Propensity |
|---|---|
| $\emptyset \to \mathrm{LacI}$ | $b_{\mathrm{LacI}} + k_{\mathrm{LacI}}/(1 + a_{2,1}[\mathrm{LacI}]^{\eta_{2,1}})$ |
| $\mathrm{LacI} \to \emptyset$ | $\gamma_{\mathrm{LacI}}[\mathrm{LacI}]$ |
| $\emptyset \to \lambda\mathrm{cI}$ | $b_{\lambda\mathrm{cI}} + k_{\lambda\mathrm{cI}}/(1 + a_{1,2}[\lambda\mathrm{cI}]^{\eta_{1,2}})$ |
| $\lambda\mathrm{cI} \to \emptyset$ | $(\gamma_{\lambda\mathrm{cI}} + \frac{0.002u(t)^2}{1250+u(t)^3})[\lambda\mathrm{cI}]$ |

Table S2: Reactions and propensities in the toggle-switch model. Parameter values are: $b_{\mathrm{LacI}} = 2.2 \times 10^{-3}$, $b_{\lambda\mathrm{cI}} = 6.8 \times 10^{-5}$, $k_{\mathrm{LacI}} = 1.7 \times 10^{-2}$, $k_{\lambda\mathrm{cI}} = 1.6 \times 10^{-2}$, $a_{2,1} = 2.6 \times 10^{-3}$, $a_{1,2} = 6.1 \times 10^{-3}$, $n_{2,1} = 3$, $n_{1,2} = 2.1$, $\gamma_{\mathrm{LacI}} = 3.8 \times 10^{-4}$, $\gamma_{\lambda\mathrm{cI}} = 3.8 \times 10^{-4}$. The UV pulse is given by $u(t) := 10 \cdot H\,(7200 - t)$ J$/m^2$ where $t$ is time (in second) after introducing UV, and $H(x)$ is the step function that takes value of one when $x \geq 0$ and zero otherwise. This formulation implies that UV pulse is turned off after a duration of two hours.

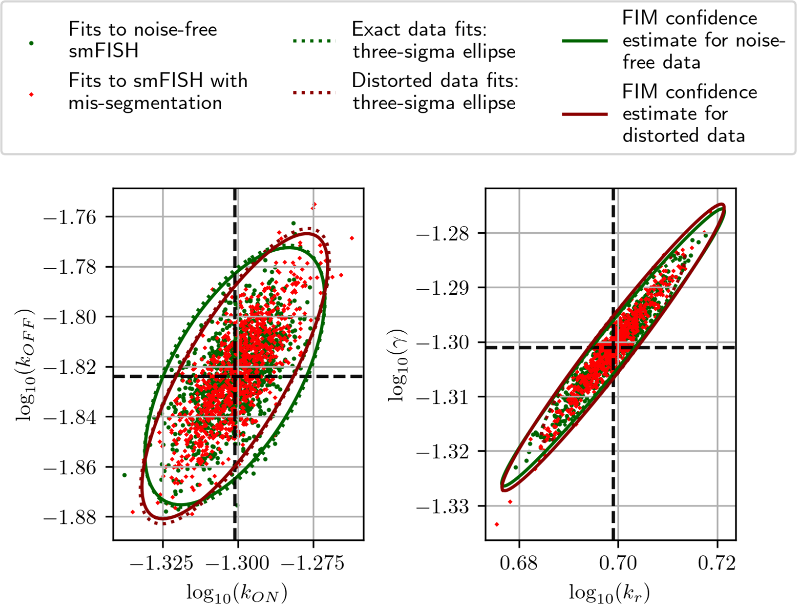


Figure S5: MLE validation for FIM-based uncertainty quantification for datasets distorted by cell segmentation noise. In the distortion simulation, we set the segmentation noise level to $\rho = 1.0$. Note how the ellipses are shrink or stretch in different directions with (red) or without (green) cell aggregations.

### S4 Two-species toggle-switch case study: Optimal combinations of different measurement modalities can provide tremendous reductions to parameter uncertainties

A common difficulty for fluorescence microscopy is the limit in the number of distinctly observable fluorophores. Most fluorescence microscopy efforts are limited to the collection of only two to four emission wavelengths, and cell segmentation often requires that one or more of those colors must be used to visualize cell nucleus or cytoplasm. It is interesting to ask how necessary it is to be able to resolve multiple mRNA species in the same cell at the same time, or is it possible to do as well with just one label at a time and in different cells. While individual partial observations will certainly incur a loss in information (and an increase in uncertainty), partial observations for different species (or combinations of species) with complementary distortion characteristics can be combined. Such "wisdom of the *diverse* crowd" can sometimes drastically improve information [6]. We revisit the toggle switch example in [3], which is based on experiments originally reported in [8] and a discrete stochastic model first formulated in [15]. The model (Fig. 4A) consists of two species, LacI and $\lambda$cI, each of which represses the production of the other (see Table S3 for reactions and propensities). Populations of isogenic cells are assumed to start from the equilibrium distribution of the toggle-switch model with no UV signal. Then a constant pulse of UV is applied to the cells with constant intensity $u := 10$ Joules/$m^2$ for two hours. We want to find the optimal time (after the initiation of the UV pulse) to collect a batch of cells from which to estimate model parameters. We explore the situation where microscopy or labeling concerns allow for fluorescent probes to label and visualize for each cell either species LacI or species $\lambda$cI, but not both simultaneously. In other words, we assume that each cell measurement can only report the abundance of a single chemical species within it. Let $\boldsymbol{C}_{\text{LacI}}$ be the probabilistic distortion operator for LacI-only measurements. We can easily see that

$$\boldsymbol{C}_{\text{LacI}}(z; (x_{\text{LacI}}, x_{\lambda\text{cI}})) = \delta_{x_{\text{LacI}}}(z),$$

where $(x_{\text{LacI}}, x_{\lambda\text{cI}})$ is the full vector of molecular counts of both species LacI and $\lambda$cI. In other words, the application of $\boldsymbol{C}_{\text{LacI}}$ on the full joint distribution of both species $\boldsymbol{p}(t)$ at time $t$ will result in the marginal distribution $\boldsymbol{p}_{\text{LacI}}(t) = \boldsymbol{C}_{\text{LacI}}\boldsymbol{p}(t)$ of species LacI at time $t$. Similarly, we can formulate the probabilistic distortion operator $\boldsymbol{C}_{\lambda\text{cI}}$ of the $\lambda$cI-only measurements, which is equivalent to the marginalization of the joint distribution $\boldsymbol{p}(t)$ into the distribution $\boldsymbol{p}_{\lambda\text{cI}}(t)$ of species $\lambda$cI. Using the Fisher Information Matrix (FIM) computation introduced above, we can compute the FIM associated with a LacI-only measurement at time $t$, which we denote by $\boldsymbol{F}_{\text{LacI}}(t)$ and similarly the single-measurement FIM $\boldsymbol{F}_{\lambda\text{cI}}(t)$ for a $\lambda$cI-only measurement.

We compute the FIM associated with experiment designs where a batch of cells is collected at a time $t$, in which $n_{\text{LacI}}$ single-cell partial observations are made for counting LacI and $n_{\lambda\text{cI}}$ cells are measured for $\lambda$cI. Assuming that the expression within each *separately measured* cell is independent from the others, the FIM associated with the experiment design outlined above is given by

$$\boldsymbol{F}(n_{\text{LacI}}, n_{\lambda\text{cI}}, t) = n_{\text{LacI}}\boldsymbol{F}_{\text{LacI}}(t) + n_{\lambda\text{cI}}\boldsymbol{F}_{\lambda\text{cI}}(t).$$

Fig. S6 compares the determinant of the FIM associated with using different combination schemes, including extreme schemes where only LacI ($n_{\text{LacI}} = 1000$, $n_{\lambda\text{cI}} = 0$) or $S_{\lambda\text{cI}}$ ($n_{\text{LacI}} = 0$, $n_{\lambda\text{cI}} = 1000$) are measured and half-and-half mixtures with $n_{\text{LacI}} = n_{\lambda\text{cI}} = 500$ and $n_{\text{LacI}} = n_{\lambda\text{cI}} = 1000$. The computed optimal sampling times associated with these combination schemes and their relative informational values are visualized in Fig. S6B (also see Supplemental Table S4 for precise numerical values). We also display the result of joint optimization of both the composition of single-species observations (constrained to not exceed $1,000$) and sampling time (Fig. S6F, also see Supplemental Text S4.1 for details). The ideal case where $1,000$ joint observations can be made is also displayed for comparison.

The advantage of combining measurements from different probe designs is profound. Indeed, when we constrain the maximum number of observations (whether partial or full two-dimensional) to $1,000$ cells, the combined measurement approach of half-and-half mixture (or its slightly more fine-tuned composition by the joint optimization displayed in Fig. S6F) results in several orders of magnitude larger determinant for the FIM than using either measurement method alone (Figs. S6B and S6C). The FIM-estimated confidence ellipse associated with the mixture design is on par with that from the full joint observations, while those associated with using only one measurement type are many orders of magnitude larger (Figs. S6D and S6E).

### S5 Exploring the effects of partial or lumped observations for the identification of spatially-compartmentalized gene expression models

We consider the model of MAPK-activated expression of the STL1 gene in yeast studied in [12] (Table S3). This model extends the telegraph model discussed in the main text in the following ways: the gene can have four states, and mRNAs are created first in the nucleus but can then be shuttled to the cytoplasm. In addition, the rate for which STL1 gene turns from the first activated state to the deactivated state is dependent on the time-varying external signal and is given by

$$k_{10}(t) = \max\left(0, k_{10}^{(a)} - k_{10}^{(b)} \cdot S(t)\right).$$

Following [12], the signal intensity $S(t)$ is given by

$$S(t) = A_{\text{hog}} \left(\frac{u(t)}{1 + u(t)/M_{\text{hog}}}\right)^{\eta},$$

where $u(t) := (1 - e^{-r_1 t})e^{-r_2 t}$ and $A_{\text{hog}}, M_{\text{hog}}, \eta, r_1, r_2$ are parameters for Hog1p concentration that we assume to be known prior to doing single-cell experiments (see Table S4 for parameter values). Our concern is to recover the gene transition rates, nuclear mRNA synthesis rates, mRNA transport rate, and mRNA degradation rates (Table S5) through joint or partial observations of mRNA copy numbers. We consider the following types of single-cell observations:

1. Joint gene-RNA: the gene state, the copy numbers of nuclear and cytoplasmic mRNAs are jointly observed per cell. This is an idealistic type single-cell observation that represents the highest amount of information that we can get from distributions of single-cell, single-molecule copy numbers.

2. Joint RNA: the gene is not observable, only the joint copy numbers of mRNAs in nucleus and cytoplasm are observed. This is the measurement type used in [12].

3. Total RNA: the total number of RNAs in both nuclear and cytoplasm is observed per cell, but the spatial information is lost.

4. Nuclear RNA-only: only nuclear RNAs are observable.

We consider an experiment in which cells are initially distributed according the stationary solution of the CME without Hog1p signal (where $k_{10}^{(b)}$ is set to zero). After osmotic shock, Hog1p relocates to the nucleus and activates the expression of STL1. Cells are collected in independent batches, of one thousand cells each, at five equally-spaced time points $\Delta t, 2\Delta t, 3\Delta t, 4\Delta t, 5\Delta t$ after applying osmotic shock. Fig. S7A shows the determinant of the FIM associated with different measurement methods and values of the sampling period $\Delta t$. The plot quantitatively confirms our intuition that the more spatial information and species we can include into a single-cell observation, the more information we can obtain from the experiment. These

| Parameter | Value |
|---|---|
| $A_{\text{hog}}$ | 9.3E9 |
| $M_{\text{hog}}$ | 2.2E-2 |
| $r_1$ | 6.1E-3 |
| $r_2$ | 6.9E-3 |
| $\eta$ | 5.9 |

Table S4: Parameters for Hog1p signal concentration, taken from [12].

different measurement methods result in different shapes and volumes of the uncertainty ellipses (Figs. S7C). Because the Nuclear RNA method discards information about the cytoplasmic mRNAs, it is unable to resolve the values of the transport and degradation rates of nuclear mRNA as seen in Fig. S7C.

Despite the great loss of information from using nuclear mRNA measurements alone, it could be potentially useful to combine it with the total mRNA measurements to obtain experiments that are more informative than those using either measurement methods alone. We consider optimizing the composition of nuclear-only measurements and total measurements at all measurement times. For simplicity, we consider only the case where the same composition is used for all measurement times. Fig. S8 shows the determinant of the FIM associated with different compositions and sampling period. It is interesting to notice that the optimal composition comes at about 833 cells total RNA measurements and 167 cells with nuclear RNA-only measurements. Despite the total mRNA being vastly more informative than nuclear-only measurements when used alone, they can be complemented by a judicious number of the less informative measurements and drastically improve information (Table S6). The uncertainties from the combined measurements are comparable to the ideal joint measurements (Fig. S10). However, in terms of the cost of imaging, nuclear mRNAs can take much less time to identify and count than the total mRNAs. For example, it is much easier for an image processing algorithm to identify and segment cell DAPI-stained nuclear regions than it is to segment the boundaries between cells [7].

| Reaction | Propensity |
|---|---|
| $G_0 \to G_1$ | $k_{01}.G_0$ |
| $G_1 \to G_0$ | $k_{10}(t).G_1$, where $k_{10}(t) = \max\left(0, k_{10}^{(a)} - k_{10}^{(b)}.S(t)\right)$ |
| $G_1 \to G_2$ | $k_{12}.G_1$ |
| $G_2 \to G_1$ | $k_{21}.G_2$ |
| $G_2 \to G_3$ | $k_{23}.G_2$ |
| $G_3 \to G_2$ | $k_{32}.G_3$ |
| $G_0 \to G_0 + \text{RNA}_{\text{nuc}}$ | $\alpha_0.G_0$ |
| $G_1 \to G_1 + \text{RNA}_{\text{nuc}}$ | $\alpha_1.G_1$ |
| $G_2 \to G_2 + \text{RNA}_{\text{nuc}}$ | $\alpha_2.G_2$ |
| $G_3 \to G_3 + \text{RNA}_{\text{nuc}}$ | $\alpha_3.G_3$ |
| $\text{RNA}_{\text{nuc}} \to \emptyset$ | $\gamma_{\text{nuc}}.\text{RNA}_{\text{nuc}}$ |
| $\text{RNA}_{\text{nuc}} \to \text{RNA}_{\text{cyt}}$ | $k_{\text{transport}}.\text{RNA}_{\text{nuc}}$ |
| $\text{RNA}_{\text{cyt}} \to \emptyset$ | $\gamma_{\text{cyt}}.\text{RNA}_{\text{cyt}}$ |

Table S3: Reactions and propensities in the spatial four-state gene expression model.

| Design | Optimal sampling period (minute) | D-optimality |
|---|---|---|
| Joint (gene, nuclear, cytoplasmic) | 3 | 2.85e+84 |
| Joint (nuclear, cytoplasmic) | 4 | 7.02e+56 |
| Nuclear RNA | 4 | 1.89e+21 |
| Cytoplasmic RNA | 4 | 4.47e+49 |
| Total RNA | 4 | 4.72e+49 |
| 833 total RNA, 167 nuclear RNA (optimal) | 3 | 1.24e+53 |

Table S6: STL1 spatial gene expression example. Optimal sampling period associated with different measurement methods (and their combinations) based on D-optimality criteria. Every design contains a total of 1,000 cels per time point.

| Parameter | Value |
|---|---|
| $k_{01}$ | `2.6E-3` |
| $k_{10}^{(a)}$ | `1.9E01` |
| $k_{10}^{(b)}$ | `3.2E04` |
| $k_{12}$ | `7.63E-3` |
| $k_{21}$ | `1.2E-2` |
| $k_{23}$ | `4.0E-3` |
| $k_{32}$ | `3.1E-3` |
| $\alpha_0$ | `5.9E-4` |
| $\alpha_1$ | `1.7E-1` |
| $\alpha_2$ | `1.0` |
| $\alpha_3$ | `3E-2` |
| $k_{\text{transport}}$ | `2.6e-1` |
| $\gamma_{\text{nuc}}$ | `2.2e-6` |
| $\gamma_{\text{cyt}}$ | `8.3e-3` |

Table S5: Parameters in the spatial four-state gene expression model. These values are taken from [12], where they were fit to the expression of STL1 gene in yeast in response to osmotic shock.

[10] Lisi Meri. Some remarks on the Cantor pairing function. *Le Matematiche*, 62, December 2007.

[11] Brian Munsky and Mustafa Khammash. The finite state projection algorithm for the solution of the chemical master equation. *The Journal of Chemical Physics*, 124(4):044104, January 2006.

[12] Brian Munsky, Guoliang Li, Zachary R. Fox, Douglas P. Shepherd, and Gregor Neuert. Distribution shapes govern the discovery of predictive models for gene regulation. *Proceedings of the National Academy of Sciences*, 115(29):7533–7538, July 2018. Publisher: National Academy of Sciences Section: Biological Sciences.

[13] J. Pfanzagl and R. Hamböker. *Parametric Statistical Theory*. De Gruyter Textbook. W. de Gruyter, Berlin ; New York, 1994.

[14] Arjun Raj and Alexander van Oudenaarden. Single-Molecule Approaches to Stochastic Gene Expression. *Annual Review of Biophysics*, 38(1):255–270, 2009.

[15] T. Tian and K. Burrage. Stochastic models for regulatory networks of the genetic toggle switch. *Proceedings of the National Academy of Sciences*, 103(22):8372–8377, May 2006.

[16] Huy Vo and Brian Munsky. A parallel implementation of the finite state projection algorithm for the solution of the chemical master equation. *bioRxiv : the preprint server for biology*, 2020.

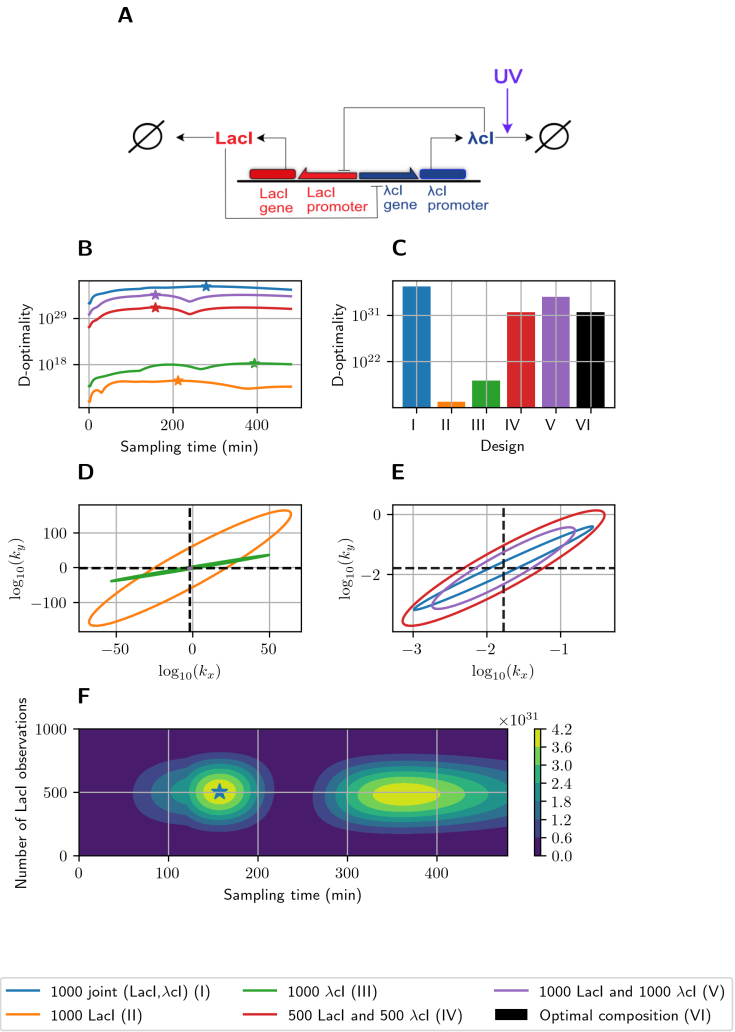


Figure S6: **Optimizing the mixture of different measurement modalities for toggle-switch model.** **(A)**: Schematic of the toggle switch model. **(B)**: D-optimality across sampling time for experiments that use different compositions of partial observations. **(C)**: D-optimality versus sampling times for the different compositions of measurements. **(D)**: FIM-estimated confidence ellipses on the $k_{\mathrm{LacI}}$-$k_{\lambda\mathrm{cI}}$ plane in log10 space associated with the composition schemes. **(E)** displays a zooming in of panel (D) to reveal the ellipses for the 1000 joint measurements scheme and the combination schemes where both LacI reporters and $\lambda$cI reporters are used. **(F)** FIM D-optimality landscape for different combinations of LacI and $\lambda$cI observation counts and sampling times (we constraint the numbers of LacI and $\lambda$cI observations to sum to 1000 and thus only show the number of LacI observations on the vertical axis). The $\star$ symbol marks the optimal solution of 502 LacI and 498 $\lambda$cI observations collected at 147 minutes.

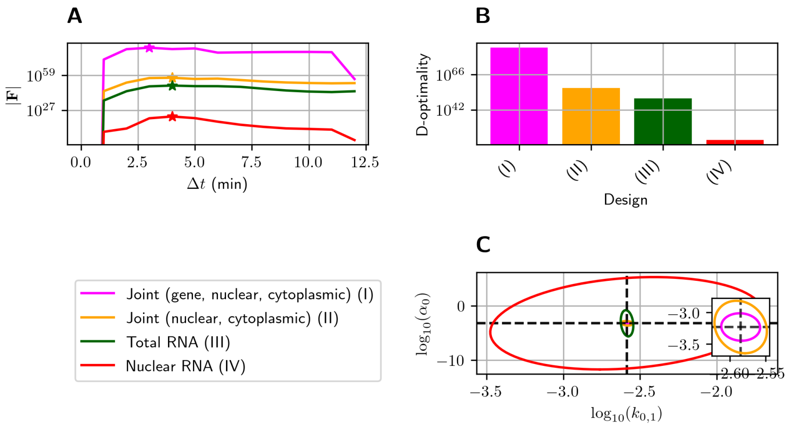


Figure S7: Information and parameter uncertainty in experiments with full and partial observations of the MAPK-activated STL1 gene expression model.

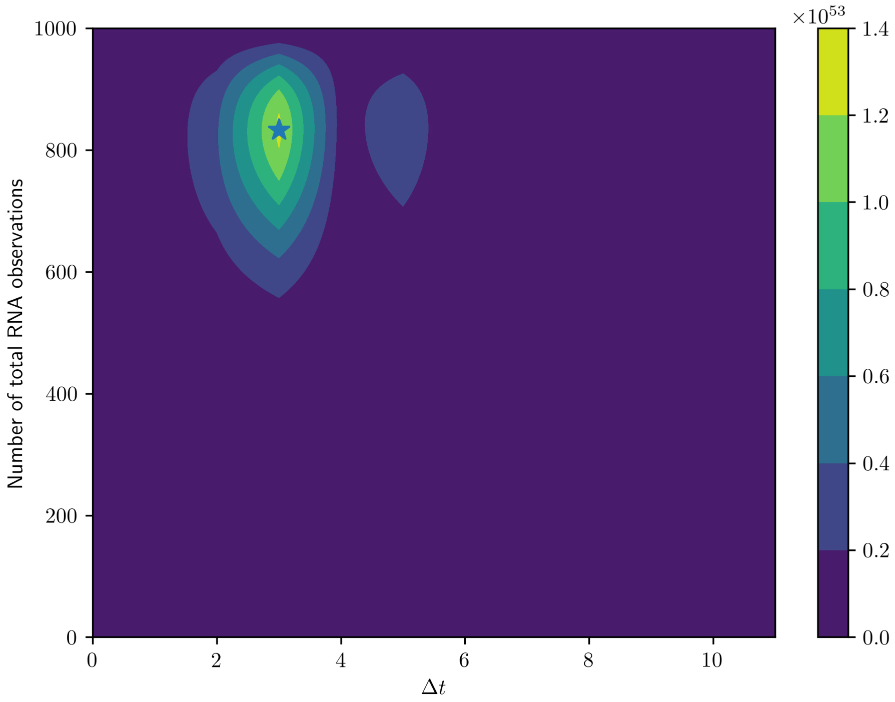


Figure S8: Determinant of the FIM associated with composite measurements for the MAPK-activated STL1 gene expression model at different sampling time periods $\Delta t$ and number of total RNA measurements per batch.

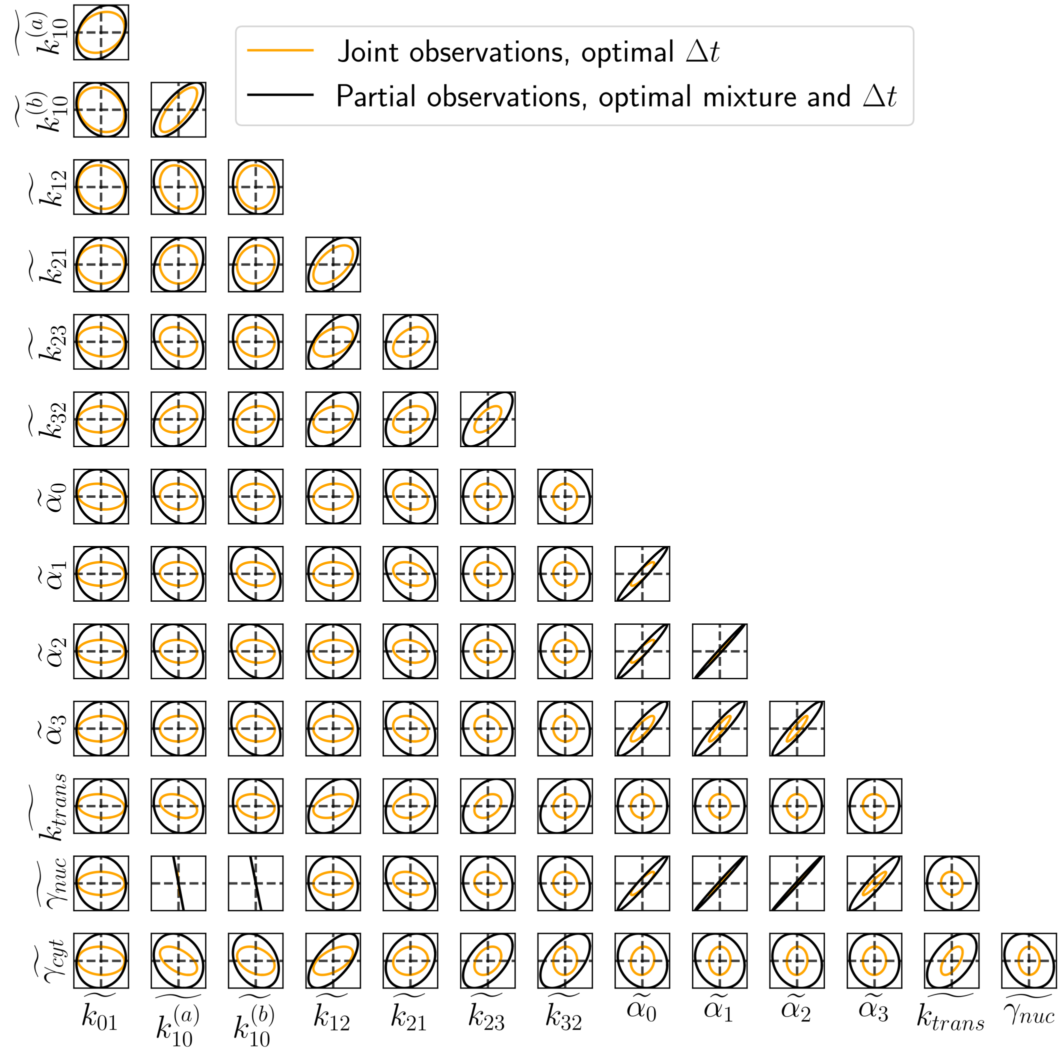


Figure S9: FIM-estimated confidence ellipses in parameter space at the optimal sampling period for each measurement types for the MAPK-activated STL1 gene expression model. In the axis labels, $\tilde{\theta}$ stands for $\log_{10}(\theta)$.

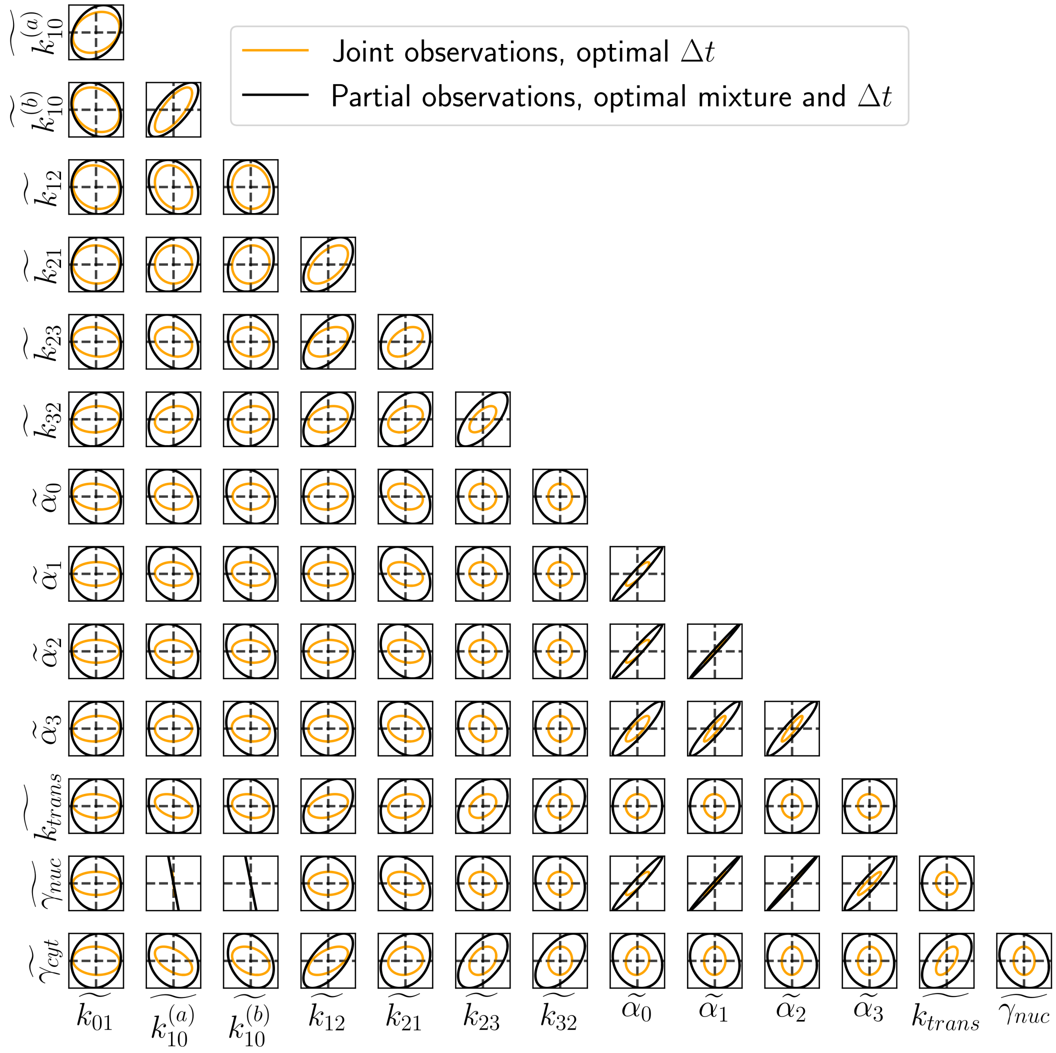


Figure S10: FIM-estimated confidence ellipses in parameter space at the optimal sampling period for each measurement types for the MAPK-activated STL1 gene expression model. In the axis labels, $\tilde{\theta}$ stands for $\log_{10}(\theta)$.

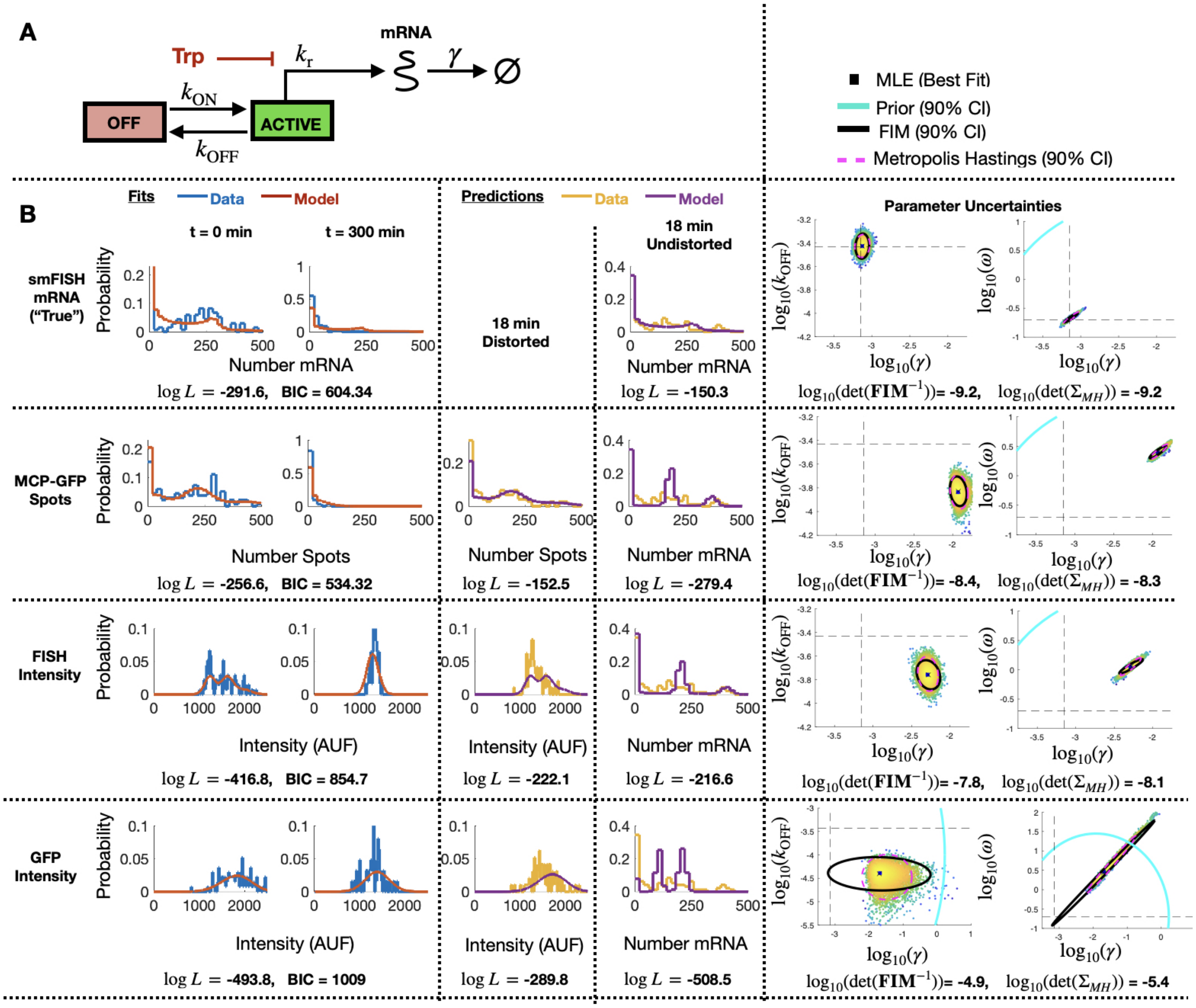


Figure S11: **Identification of Simplified Stochastic Model.** **(A)** Schematic of the simplified 2-state gene expression model. **(B)** Results for model fitting, prediction, and uncertainty quantification for measurements based smiFISH spots (top row), MCP-GFP spots (row 2), total FISH intensities (row 3) and GFP intensities (row 4). Left two columns show the measured and model-fitted probability mass vectors (PMV) at 0 and 300 min after 5$\mu$M Tpt. Third column shows the model-predicted and measured PMV for the corresponding (distorted) measurement modality at 18 min after 5$\mu$M Tpt. Fourth column shows the model prediction without distortion and measured PMV for the smFISH mRNA count at 18 min after 5$\mu$M Tpt. All histograms use a bin size of 20. Log-likelihood values for all model-data comparisons (and BIC values for fitting cases, $k$ = 4 parameters, N = 197 cells) are computed without binning and are shown below the corresponding histograms. Right two columns show joint parameter uncertainty for model estimation using data for 0 and 300 min after 5$\mu$M Tpt. In each case, the 90% CI for prior is shown in cyan; Metropolis Hastings samples (N = 20,000) are shown in dots; 90% CI for posterior is shown in dashed magenta; and FIM-based estimate of 90% CI is shown in black. Horizontal and vertical dashed black lines denote the "true" parameters and are defined as the MLE when using fit to the smFISH counts and using all time points. Determinant of inverse FIM and covariance of MH samples is shown below each pair of uncertainty plots (both use log base 10).

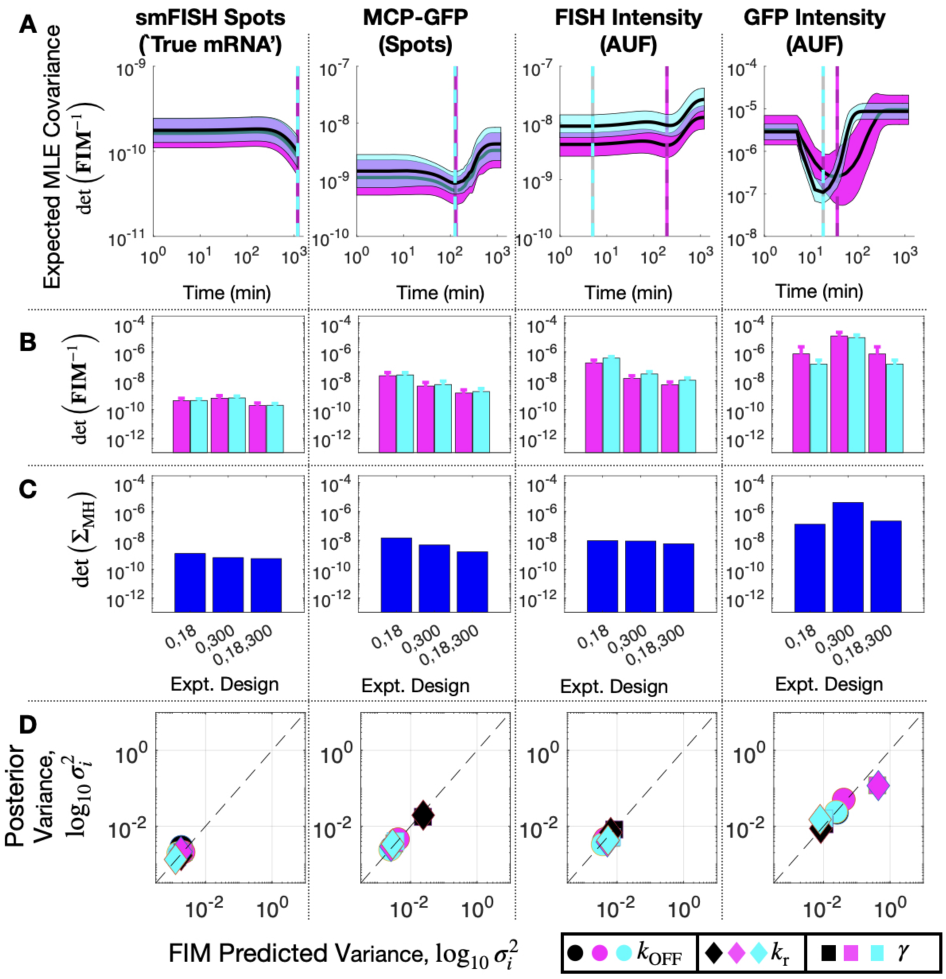


Figure S12: **Design of Subsequent Experiment for the Simplified Model. (A)** Expected volume of uncertainty ($\det(\mathbf{FIM}^{-1})$) versus time of third measurement assuming 100 cells and measurement of: (left to right) smiFISH mRNA, MCP-GFP spots, FISH intensity, or GFP intensity. Solid lines and shading denote mean $\pm$ SD for 20 parameter sets selected from MH chains after fitting initial data (magenta, $t$ =(0,300) min) or final data (cyan, $t$ =(0,18,300) min). Cyan and magenta vertical lines denote the optimal design for the third experiment time assuming the corresponding parameter values. **(B)** Expected volume of MLE uncertainty ($\det(\mathbf{FIM}^{-1})$) for different sets of experiment times and measurement modalities and averaged over 20 parameters sets sampled from the MH chains for initial fit (magenta) or final parameter estimates (cyan). **(C)** Volume of MLE uncertainty ($\det(\Sigma_{\mathrm{MH}})$ estimated from MH analysis in the same experiment designs as **B**. **(D)** Posterior variance versus FIM prediction of variance for each parameter (symbol key at bottom right), for each measurement modality (different columns) and for analyses based on different sets of data: $t$ =(0,18) min (black), $t$ =(0,300) min (magenta), or $t$ =(0,18,300) min (cyan). All MH analyses contain 20,000 samples. Measurements include 135 cells at $t = 0$, 96 at $t = 18$, and 62 at $t = 300$. Parameter uncertainties defined in log base 10 for all panels.
